# Screening of chemical libraries for new antifungal drugs against *Aspergillus fumigatus* reveals the potential mechanism of action of miltefosine

**DOI:** 10.1101/2021.05.19.444908

**Authors:** Thaila Fernanda dos Reis, Maria Augusta Crivelente Horta, Ana Cristina Colabardini, Caroline Mota Fernandes, Lilian Pereira Silva, Rafael Wesley Bastos, Maria Vitória de Lazari Fonseca, Fang Wang, Celso Martins, Márcio L. Rodrigues, Cristina Silva Pereira, Maurizio Del Poeta, Koon Ho Wong, Gustavo H. Goldman

**Affiliations:** Faculdade de Ciências Farmacêuticas de Ribeirão Preto, Universidade de São Paulo, Ribeirão Preto, Brazil; MicroControl Innovation Ltda, Ribeirão Preto, São Paulo, Brazil; Department of Microbiology and Immunology, Stony Brook University, Stony Brook, New York, USA; Faculty of Health Sciences, University of Macau, Macau SAR of China; Instituto de Tecnologia Química e Biológica António Xavier, Universidade Nova de Lisboa (ITQB NOVA), Oeiras, Portugal; Instituto Carlos Chagas (ICC), Fundação Oswaldo Cruz – Fiocruz, Curitiba/PR, Brazil; Instituto de Microbiologia Paulo de Góes, Universidade Federal do Rio de Janeiro (UFRJ), Rio de Janeiro/RJ, Brazil; Veteran Administration Medical Center, Northport, New York, USA; MicroRid Technologies Inc., 86 Deer Park Road, Dix Hills, New York, USA; Division of Infectious Diseases, School of Medicine, Stony Brook University, New York, USA; Institute of Translational Medicine, Faculty of Health Sciences, University of Macau, Avenida da Universidade, Taipa, Macau SAR of China; MoE Frontiers Science Center for Precision Oncology, University of Macau

**Author notes:** Corresponding author: Dr. Gustavo H. Goldman, Departamento de Ciências Farmacêuticas, Faculdade de Ciências Farmacêuticas de Ribeirão Preto, Universidade de São Paulo, Av. do Café S/N, CEP 14040-903, Ribeirão Preto, São Paulo, Brazil, /81.

**Keywords:** *Aspergillus fumigatus*, drug repurposing, miltefosine, sphingolipids, transcription factor

## Abstract

*Aspergillus fumigatus* is an important fungal pathogen and the main etiological agent of aspergillosis, a disease characterized by a noninvasive process that can evolve to a more severe clinical manifestation called invasive pulmonary aspergillosis (IPA) in immunocompromised patients. The antifungal arsenal to threat aspergillosis is very restricted. Azoles are the main therapeutic approach to control IPA, but the emergence of azole-resistant *A. fumigatus* isolates has significantly increased over the last decades. Therefore, new strategies are necessary to combat aspergillosis and drug repurposing has emerged as an efficient and alternative approach for identifying new antifungal drugs. Here, we used a screening approach to analyze *A. fumigatus* in vitro susceptibility to 1,127 compounds. *A. fumigatus* was more susceptible to 10 compounds, including miltefosine, a drug that displayed fungicidal activity against *A. fumigatus*. By screening an *A. fumigatus* transcription factor null library, we identified a single mutant, which has the *rmiA* (resistant to miltefosine) gene deleted, conferring a phenotype of susceptibility to miltefosine. The transcriptional profiling (RNA-seq) of the wild-type and the Δ*rmiA* strains and the Chromatin Immunoprecipitation coupled to next generation sequencing (ChIP-Seq) of a RmiA-tagged strain exposed to miltefosine revealed genes of the sphingolipids pathway that are directly or indirectly regulated by RmiA. Sphingolipids analysis demonstrated that the mutant has overall decreased levels of sphingolipids when growing in the presence of miltefosine. The identification of RmiA represents the first genetic element described and characterized which plays a direct role in miltefosine response in fungi.

**Author summary:** The filamentous fungus *Aspergillus fumigatus* causes a group of diseases named aspergillosis and their development occurs after the inhalation of conidia dispersed in the environment. Very few classes of antifungal drugs are available for aspergillosis treatment, *e.g.*, azoles, but the emergence of global resistance to azoles in *A. fumigatus* clinical isolates has increased over the last decades. Repositioning or repurposing drugs already available on the market is an interesting and faster opportunity for the identification of novel antifungals agents. By using a repurposing strategy, we identified 10 different compounds that impact A. *fumigatus* survival. One of these compounds, miltefosine, demonstrated fungicidal activity against *A. fumigatus*. The mechanism of action of miltefosine is unknown and aiming to get more insights about it, we identified a transcription factor RmiA (<u>R</u>esistant to <u>mi</u>ltefosine) important for miltefosine resistance. Our results suggest that miltefosine plays antifungal activity against *A. fumigatus* interfering in the sphingolipids biosynthesis.

## Introduction

Fungi are widespread in nature surviving as saprophytic organisms or associated with animals and plants where they can behave as commensal or opportunistic organisms. In humans, pathogenic fungi can cause both superficial and invasive infections giving rise to the death of millions of people annually (1–3). *Cryptococcus, Candida, Aspergillus*, and *Pneumocystis spp* are responsible for the most representative invasive fungal infections (1) showing death rates as high as tuberculosis and malaria (2, 4, 5). The levels of mortality are dependent on the host immune system integrity, being particularly important for immunocompromised patients (6–8). These individuals comprise a risk group that is expanding quickly due to the increasing number of immune-deficient patients who underwent transplant or chemotherapy, and patients under therapy with high dosage of corticosteroids (9–11).

*Aspergillus spp*. cause a group of diseases collectively named aspergillosis and their development occur after the inhalation of conidia dispersed in the environment (12). In immunocompetent patients the development of aspergillosis is mainly characterized by noninvasive diseases including aspergilloma, chronic necrotizing pulmonary aspergillosis, chronic cavitary pulmonary aspergillosis, and chronic fibrotic pulmonary aspergillosis, which are together defined as chronic pulmonary aspergillosis (12–16). These diseases present different manifestations but are in general related to a preexisting cavitary lung disease and classical features of systemic fungal disease are always absent (17, 18). Additionally, the allergic bronchopulmonary aspergillosis (ABPA) is a severe atopic disease caused by *Aspergillus*, mainly *A. fumigatus* (19, 20). The disease originates from the sensitization to fungal allergens of the host who present high levels of IgE (21). Patients with cystic fibrosis and other genetic diseases that affect the respiratory system are more predisposed to ABPA (19, 22, 23). Finally, the invasive pulmonary aspergillosis (IPA) is an important clinical manifestation caused by *Aspergillus* spp. presenting high levels of mortality in immunocompromised patients (1, 24). IPA is the most common invasive fungal infection in recipients of both hematopoietic stem cells and solid-organ transplants (1, 24). In this group of high-risk patients for IPA, *A. fumigatus* represents the major cause of the disease reaching up to 90% of mortality (9–12, 25).

Very few classes of antifungal drugs are available for IPA treatment, such as polyenes (amphotericin B), azoles (itraconazole, posaconazole, voriconazole, and isavuconazole), and echinocandins (caspofungin) (26–29). Although both amphotericin B and echinocandins can be used to treat IPA, these drugs have clinical limitations. Amphotericin B shows high levels of nephrotoxicity and side effects while echinocandins are not fully recommended as monotherapy for IPA (9, 13, 30–32). So far, the administration of triazoles is the first therapeutic approach applied to control *A. fumigatus* infections showing the most prominent usage in the medical field (13, 33). Among them, itraconazole (introduced in 1990s), voriconazole (introduced in 2002) and posaconazole (introduced in 2006) are the most common drugs utilized for the treatment of aspergillosis (34). Voriconazole is the primary treatment against IPA followed by liposomal amphotericin B (L-AMB) and echinocandins, which are recommended as a second line therapy (35) (13, 33). Moreover, the activity of isavuconazole, a new extended-spectrum triazole drug, has been recently tested against *Aspergillus* (36–39).

The number of azole-resistant *A. fumigatus* clinical isolates has dramatically increased over the last decades has become a major concern (35, 40–45). Additionally, azoles are also used in the agriculture to combat plant pathogenic fungi and recently, its usage for agricultural purposes has been linked to the emergence of azole-resistant isolates among human fungal pathogens (40, 46–49). Therefore, the emergence of global resistance to currently available antifungals agents represent a significant threat to immunosuppressed patients, as the current arsenal of antifungal drugs is very limited.

*A. fumigatus* has developed different azole resistance mechanisms, e.g. amplification of the gene encoding the drug target, Cyp51/Erg11 14-alpha demethylase; drug target modification, and/or overexpression of efflux pumps (50–54). Specifically, These events are linked to amino acid substitution(s) in the target Cyp51A protein, tandem repeat sequence insertions at the cyp51A promoter, and overexpression of the ABC transporter Cdr1B (55–57). The scenario became even more dramatic since the emergence of fungal isolates intrinsically resistant to other commercial antifungals available. For instance, Some clinical isolates of *A. flavus* and *A. terreus*, also responsible for cases of IPA, have been reported as intrinsically resistant to amphotericin B (12, 29, 58– 61).

This situation highlights the need to understand the mechanisms of drug resistance and tolerance, and the search for novel antifungal agents (62, 63). As a few antifungal compounds are coming to market because their development is time-consuming and expensive, repositioning or repurposing drugs which are already licensed is an interesting and faster opportunity for the identification of novel antifungals agents (64–66). By using the repurposing strategy, many compounds have already been identified as new potential drugs against several diseases including parasitosis, protozooses and mycoses (64, 66–71). Here, screened two compound collection to analyze *A. fumigatus in vitro* susceptibility for compounds present in two compound libraries. The first library has active compounds against neglected diseases (The Pathogen Box) while the second one includes drugs previously approved for usage against human diseases [National Institutes of Health (NIH) clinical collection (NCC)]. We showed here that *A. fumigatus* was susceptible to at least 10 different compounds from the two libraries. One of these compounds, miltefosine, a drug mainly used in the treatment of visceral and cutaneous leishmaniasis (72, 73), demonstrated fungicidal activity against *A. fumigatus*. Aiming to get more insights about the mechanism of action of miltefosine, we screened an *A. fumigatus* transcription factor null mutant library (484 null mutants) and identified a single mutant highly sensitive to miltefosine. The gene deleted in this mutant was named *rmiA* (<u>r</u>esistant to <u>mi</u>ltefosine). A combination of RNA seq transcriptome and Chromatin Immunoprecipitation (ChIP) coupled to next generation sequencing (Seq) studies revealed differentially expressed genes directly or indirectly regulated by RmiA. The sphingolipids profiling of the wild-type and the Δ*rmiA* strains exposed to miltefosine revealed that the mutant has overall lower levels of sphingolipids comparing to the wild-type. Our results suggest that miltefosine plays antifungal activity against *A. fumigatus* by directly interfering in the sphingolipids biosynthetic pathway.

## Results

### Screening of the Pathogen Box and NIH Clinical Library

In order to find known compounds that are active against *A. fumigatus*, we tested its susceptibility to two chemical drug libraries, (i) the Pathogen Box (containing 400 compounds, see https://www.mmv.org/mmv-org) and the National Institutes of Health (NIH) clinical collection (NCC) (containing 727 compounds; see https://pubchem.ncbi.nlm.nih.gov/source/NIH%20Clinical%20Collection) through Minimal Inhibitory Concentration (MIC) assays. In total, combining both libraries 1,127 compounds were assessed by using MIC values up to 25 µM. *A. fumigatus* was susceptible to four known antifungal agents present in these collections (posaconazole, difenoconazole, bitertanol and amphotericin B; MIC values 5 µM, 5 µM, 5 µM and 10 µM, respectively). These results supported the reliability of the screening approach. *A. fumigatus* was also susceptible to other compounds with MIC values ranging from 1.56 to 25 µM (Table 1). In Table 1, we describe the compound name, the MIC detected in our screening, the current usage purpose (Description) and the mode of action (if known) for the 10 compounds (Table 1). These compounds include: (i) two azole salts, econazole and oxiconazole, expected to inhibit to some extent *A. fumigatus* growth, (ii) fluvastatin, a statin drug class used for hypercholesterolemia treatment, (iii) mesoridazine, a piperidine neuroleptic drug, (iv) cisapride, a parasympathomimetic drug acting as a serotonin 5-HT_4_ agonist, (v) indinavir sulfate, a protease inhibitor used in anti-HIV cocktails, (vi) enalaprilat, an angiotensin-converting enzyme inhibitor, (vii) vincristine sulfate, an inhibitor of microtubule formation in the mitotic spindle, (viii) iodoquinol, an anti-amoebiasis agent with unknown mechanism of action and (ix) miltefosine, an anti-Leishmania compound with unknown mechanism of action (Table 1).

**Table 1.**
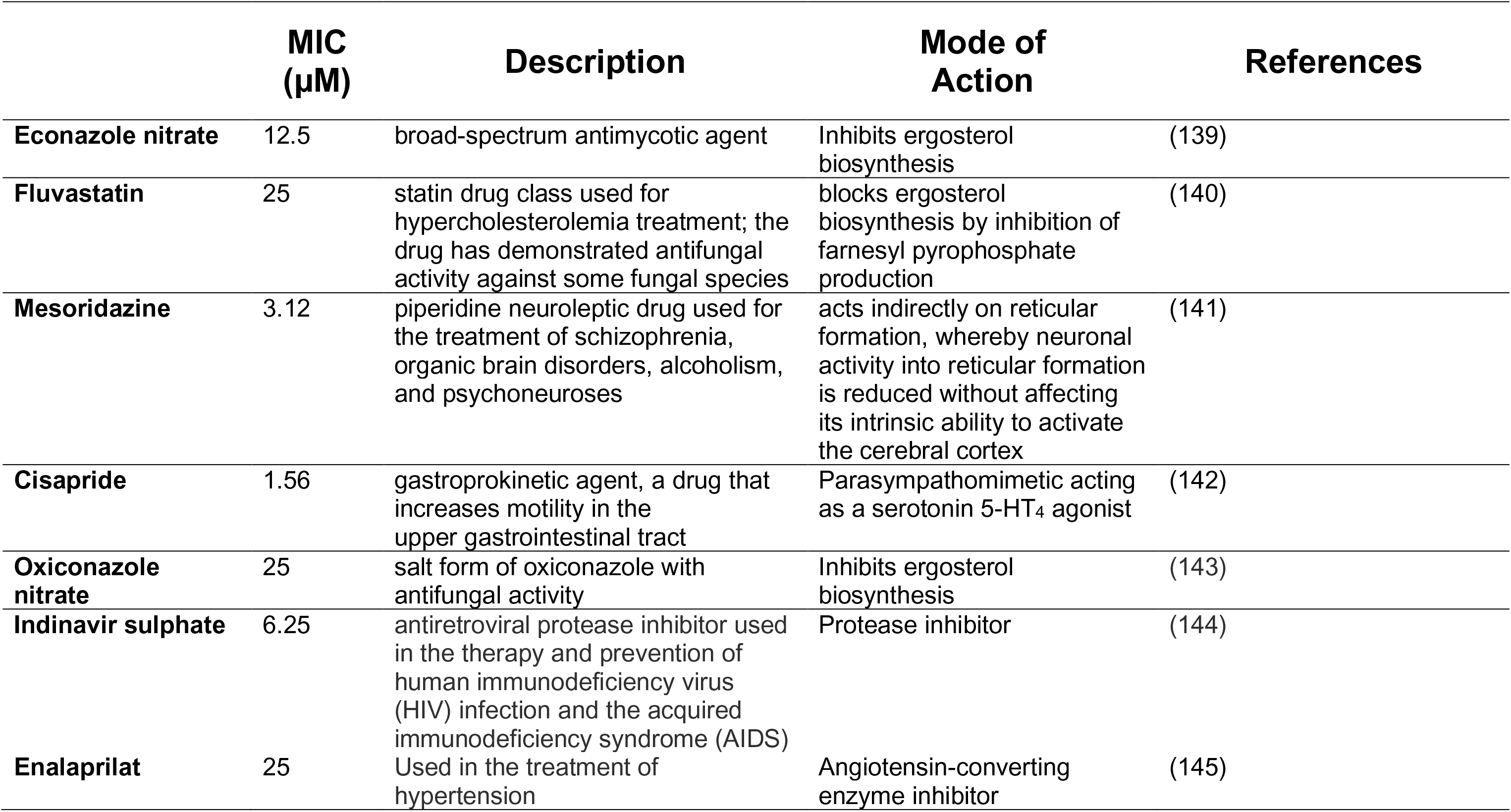
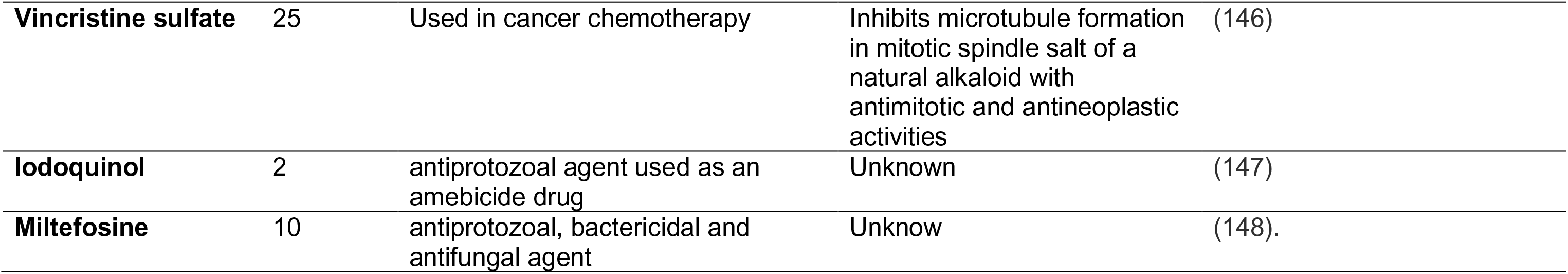
MIC values for NIH Clinical collection and Pathogen box compounds against *A. fumigatus*

To determine if these compounds are fungicidal or fungistatic, the *A. fumigatus* conidial viability was tested after 48 h of exposure to each compound at its corresponding MIC (Figure 1A). Five compounds (fluvastatin, cisapride, indinavir sulphate, vincristine sulphate, and miltefosine) had 100 % fungicidal while six had fungistatic mechanism of action with 80 to 95 % of conidial killing at MIC concentration (Figure 1A).

**Figure 1.**
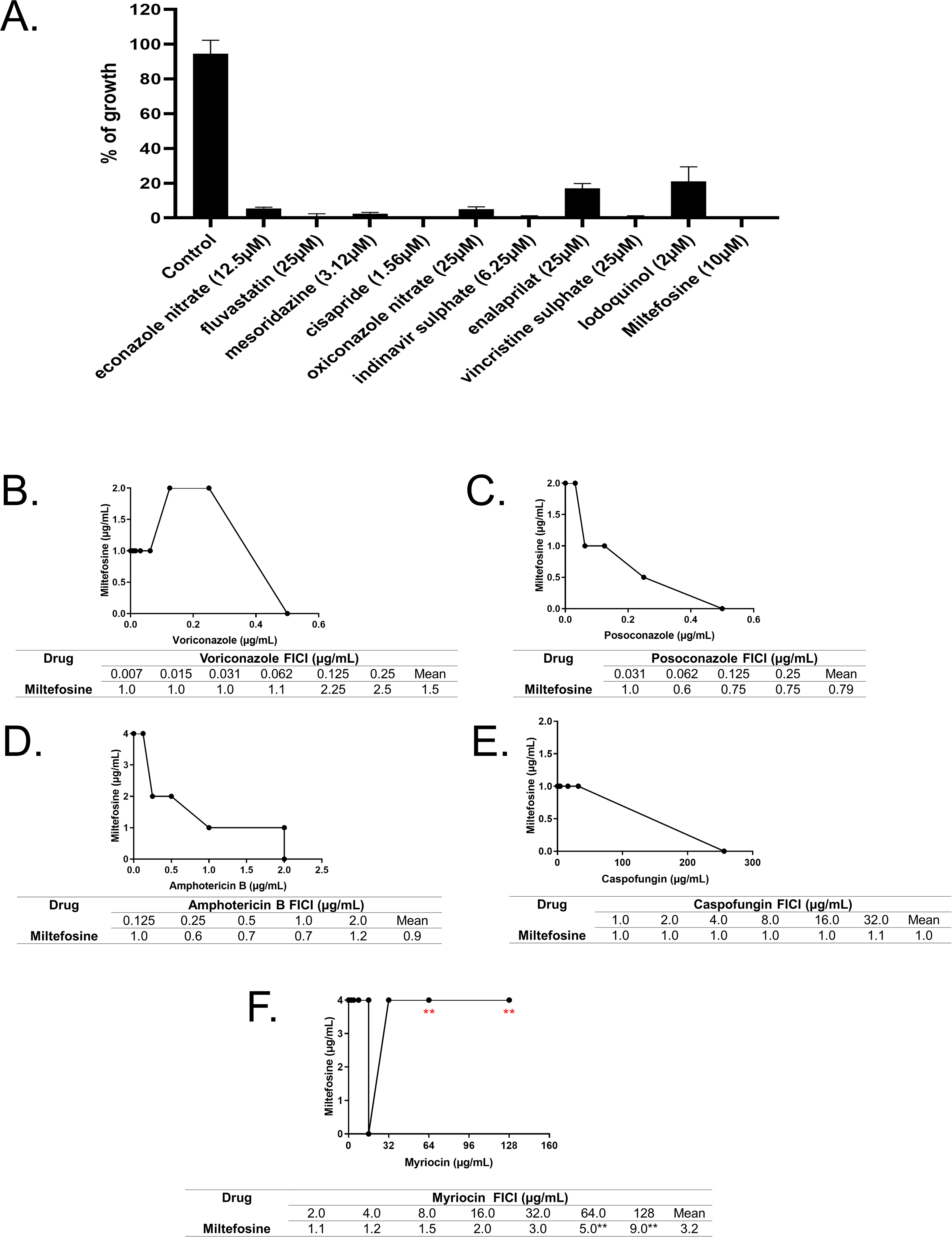
Miltefosine is a potential new anti-aspergillosis compound and shows its interaction with the sphingolipid inhibitor myriocin. (A). Screening of chemical libraries reveals potential new anti-aspergilosis compounds. (B) Interaction between miltefosine and Posaconazole. (C) Interaction between miltefosine and voriconazole. (D) Interaction between miltefosine and amphotericin B. (E) Interaction between miltefosine and caspofungin. (F) Interaction between miltefosine and myriocin

These results suggest that some of these compounds are fungicidal and can act directly in specific *A. fumigatus* cell targets while others (like cisapride and enalaprilat) could be a lead compound in antifungal drug discovery.

### Miltefosine displays antagonistic interaction with myriocin, a sphingosine biosynthesis inhibitor

We decided to investigate miltefosine in more detail because it is a fungicidal drug with an unknown mechanism of action. The combination between drugs is commonly used in clinical practice aiming to potentialize the antifungal effect of the drugs (74). Furthermore, the combination assay can help unraveling the mechanism of action of the drugs and how this may vary according to the concentration (75). To check if miltefosine has any interaction with other antifungal drugs, we combined this compound (ranging from 0.001 to 8.0 µg/ml, MIC value of 10 µM corresponds to 4 µg/ml) with different antifungal drugs (Figures 1B to 1E). Miltefosine was combined with posaconazole [0.03 - 2.0 µg/ml], voriconazole [0.0007 - 0.5 µg/ml], amphothericin B [0.06 - 4.0 µg/ml] and caspofungin [4.0 - 256.0 µg/ml]) (Figures 1B to 1E). Using the checkerboard microdilution method, the interaction between miltefosine and the other compounds was determined through the fractional inhibitory concentration index (FICI). The interaction between the drugs was classified as synergic (FICI ≤ 0.5), indifferent (0.5 < FICI ≤4.0) or antagonic (FICI > 4.0) (76). Under the assayed conditions, the FICI index varied from 1- 2.5 in the combination of voriconazole and miltefosine, 1- 0.6 between posaconazole and miltefosine, 0.6 – 1.2 between amphothericin B and miltefosine and 1.0 -1.1 between caspofungin and miltefosine. These data show that the addition of miltefosine did not affect the antifungal effects of the tested clinical antifungals against *A. fumigatus* indicating that there is no interaction between them.

There are evidences in the literature showing that miltefosine can affect the sphingolipid metabolism in trypanosomatids (77, 78). To check if miltefosine could display any interaction with drugs that affect the cellular lipids biosynthesis, we combined different concentrations of miltefosine [0.001 to 8.0 µg/ml] and myriocin [2.0 – 128 µg/ml] (Figure 1F), an inhibitor of serine palmitoyltransferase, the first step in sphingosine biosynthesis (79). At low concentrations of the drugs, indifferent interaction was observed. Interestingly, at high concentrations, myriocin impaired the antifungal effects of miltefosine against *A. fumigatus* demonstrating an antagonistic effect between these compounds (Figure 1F). Considering that myriocin has only one single target identified, this result may suggest that exist a component of the sphingolipid pathway important to the antifungal effect of miltefosine.

### RmiA is the major transcription factor that mediates miltefosine response in *A. fumigatus*

To assess if there are transcriptional programs modulating the tolerance response to miltefosine, a library of 484 *A. fumigatus* transcription factor (TF) null mutants (80) was screened for sensitivity to miltefosine [0.001 to 8.0 µg/ml]. A primary screening using 96-well plates identified six TF null mutants with different susceptibilities to miltefosine. To validate the differential susceptibility of these mutants to miltefosine, the 6 TF null mutants were grown in the absence or presence of different miltefosine concentrations, and their radial growth were measured (Figure 2). When compared to the wild-type strain, we observed discrete differences in five of these mutant strains (Figure 2). The Δ*mcnB* (AFUA_5G05600 that encodes a homologue of *A. nidulans* McnB, a multicopy supressor of *A. nidulans nimA1*, (81) showed about 20 % growth inhibition (Figures 2A and 2B) while Δ*pacC* (AFUA_3G11970 that encodes PacC, a protein important for pH regulation (82) showed about 50 % inhibition compared to the wild-type strain at 8 µg/ml (Figure 2A and 2B). The Δ*sslA* (AFUA_5G04333 that encodes a homologue of *Saccharomyces cerevisiae* Ssl1p, a subunit of the general transcription factor TFIIH) has about 50 % growth inhibition compared to the wild-type strain at 8 µg/ml (Figures 2A and 2B). The Δ*sebA* (AFUA_4G09080 encodes a TF important to cope with different kinds of stress (83) showed a 40 % inhibition to miltefosine 8 µg/ml while the wild-type is inhibited 60 % at this concentration (Figures 2A and 2B). The AFUA_5G03030 null mutant has 65 % growth inhibition compared to the wild-type strain at miltefosine 8 µg/ml (Figure 2). Notably, the AFUA_2G12070 mutant was unable to grow at 4 µg/ml miltefosine (Figures 2A, 2B, 3A and 3B). AFUA_2G12070 encodes a 492 amino acids novel fungal Zn_2_-Cys_6_ transcription factor (http://pfam.xfam.org/family/PF00172#Zn2/Cys6). We named this gene *rmiA* (*<u>r</u>*esistant to *<u>mi</u>*ltefosine). The Δ*rmiA* was complemented and the complementing strain Δ*rmiA:rmiA^+^* presented a reversible phenotype in terms of miltefosine sensitivity indicating that the miltefosine sensitivity phenotype of Δ*rmiA* is due to the specific deletion of the *rmiA* gene (Figures 3A and 3B). The Δ*rmiA* mutant has no differential susceptibility to different stressing conditions such as growth on increasing concentrations of NaCl, Calcofluor white, sorbitol, CaCl_2_ and menadione (Supplementary Figure S1).

**Figure 2.**
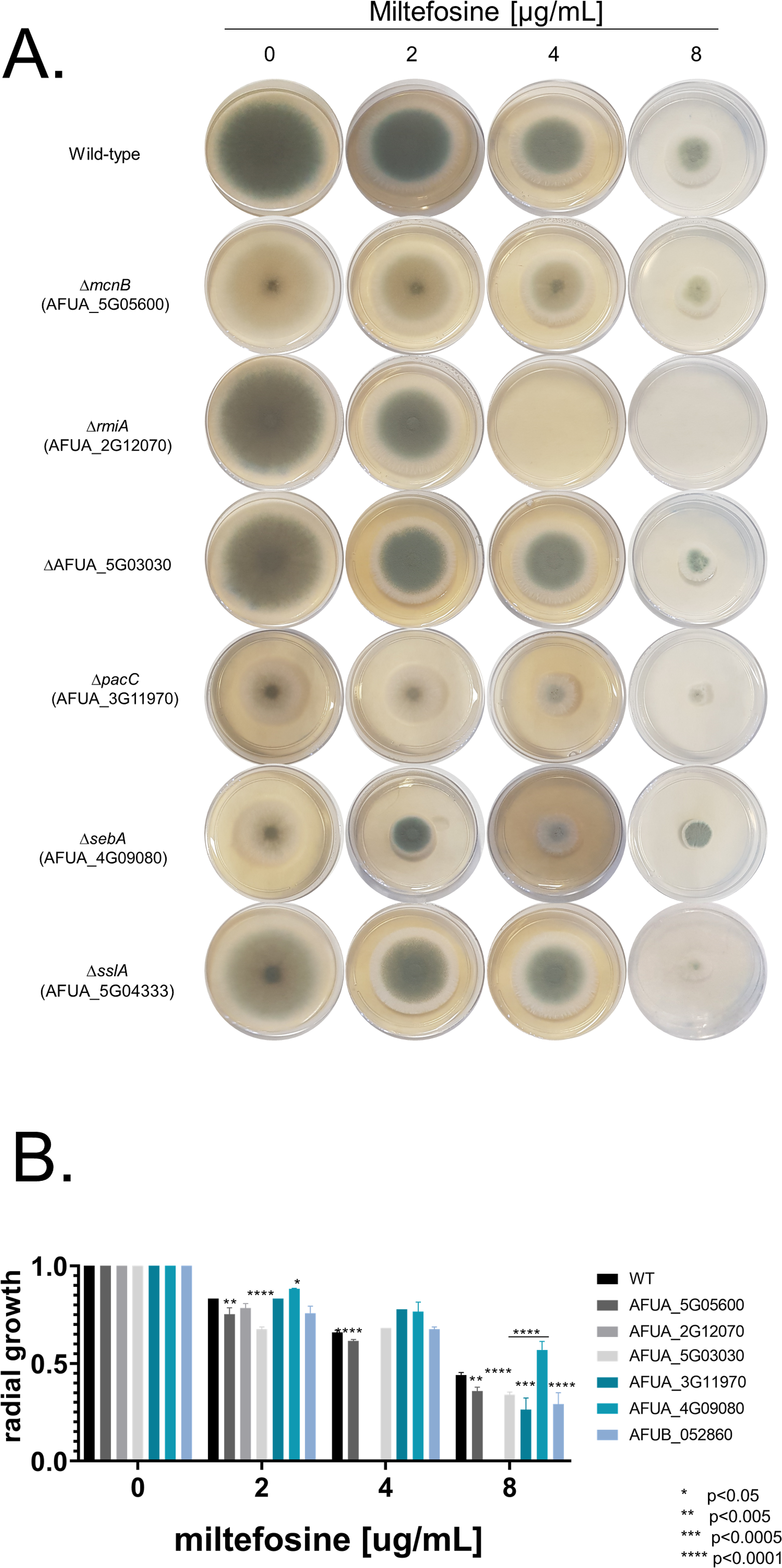
Radial growth of transcription factor (TF) null mutants in presence of miltefosine. (A) A total of 1 x 10^5^ conidia of each specie was inoculated on MM supplemented or not with increasing concentration of miltefosine. Plates were incubated for 3 days at 37°C. (B) Quantification of the results obtained in (A). For each strain three independent experiments were realized and the graphic shows the mean ± standard deviations.

**Figure 3.**
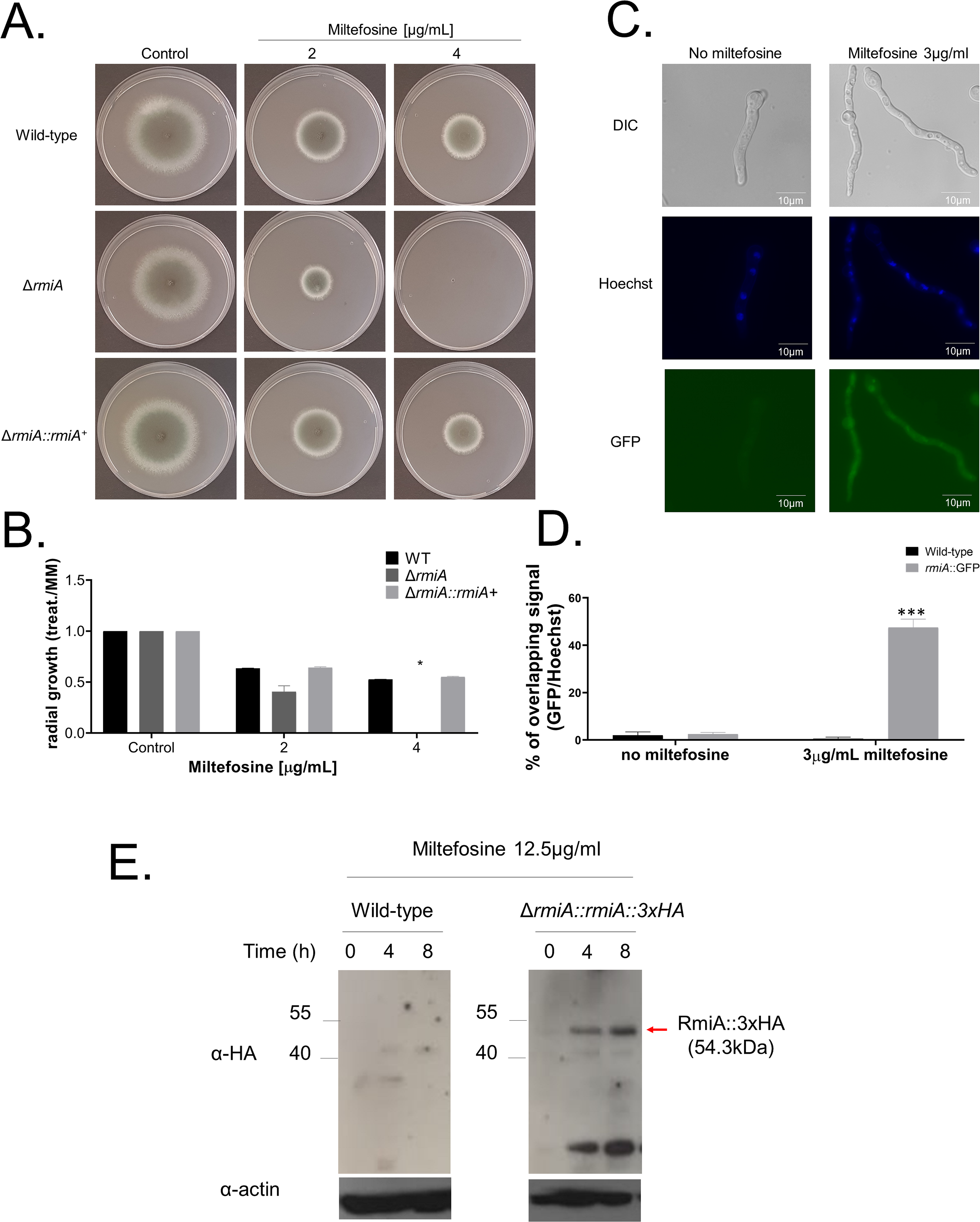
Molecular characterization of *rmiA*. (A) Growth phenotypes of the wild-type, Δ*rmiA* and Δ*rmiA::rmiA*^+^ strains grown for 3 days on solid MM supplemented with increasing concentrations of miltefosine. (B) Graphical quantification of fungal growth presented in (A). The results are the average of three repetitions ± standard deviation of three repetitions. (C) RmiA:GFP translocates to the nucleus under exposure to miltefosine. (D) Graphical quantification of RmiA:GFP location shown in (C). The results are the average of three repetitions ± standard deviation of 30 germlings for each repetition. (E) Western blotting assay showing the RmiA:HA expression after 0, 4 and 8h of incubation with 12.5 µg/mL miltefosine. Anti-HA antibody was used to detect the recombinant protein. Anti-actin antibody was used as a loading control. Statistical analysis was performed using one-tailed, paired *t* tests for comparisons to the control condition (*, p <0.05; ***,p< 0.001;)

Aiming to localize the RmiA, we constructed a functional C-terminal RmiA:GFP strain (Supplementary Figure S2) that showed no fluorescence in the absence of miltefosine (Figures 3C and 3D). However, when the RmiA:GFP strain was shifted 15 minutes to MM supplemented with 3 µg/ml of miltefosine, RmiA:GFP can be detected in about 50 % of the nuclei (Figures 3C and 3D). In addition, we also constructed a functional RmiA:3xHA strain (Supplementary Figure S2). This strain was grown in VMM and further exposed to RPMI supplemented (or not) to an inhibitory concentration of miltefosine (12.5 µg/ml) for 4 and 8 min. A very faint band of 54.3 kDa corresponding to RmiA:3xHA was observed in the control not exposed to miltefosine while increased intensity bands were observed in 4 and 8 h exposure to miltefosine (Figure 3E).

These results indicate that the RmiA protein quickly translocates to the nucleus and its expression is also increased upon miltefosine exposure.

### Miltefosine induces necrosis-like cell death and increases mitochondrial fragmentation in *A. fumigatus*

*A. fumigatus* form mitochondrial tubular and highly dynamic networks that are fragmented in the presence of antifungal and oxidative stressing agents such as hydrogen peroxide (84, 85). This increased mitochondrial fragmentation has been associated as a marker for cell death (85). Propidium Iodide (PI) is a fluorescent DNA-binding dye that freely penetrates cell membranes of dead or dying cells, but is excluded from viable cells. Late apoptosis and early necrosis is characterized by an increased number of PI-positive cells. To evaluate the effects of miltefosine and PI on the mitochondrial morphology and viability, germlings from the wild-type, Δ*rmiA* and Δ*rmiA::rmiA^+^* strains were treated with 3 µg/ml of the drug for 0, 5 or 10 min and green MitoTracker (a mitochondrial fluorescent probe) or PI were added and further analyzed by fluorescent microscopy (Figure 4A). In the absence of miltefosine an intact mitochondrial network was observed in all the three strains. However, upon 5 minutes of miltefosine exposure, the *ΔrmiA* cells showed about 60% of mitochondrial fragmentation evidenced by the presence of a punctuated fluorescent pattern observed in the cytoplasm of the cells (Figures 4A and 4B), while in the wild-type and complemented strains the levels of mitochondrial fragmentation were much lower, 20 and 30 %, respectively (Figures 4A and 4B). When the wild-type and the Δ*rmiA::rmiA+* germlings were left unexposed to miltefosine, about 5% of cells were stained by PI while in the Δ*rmiA* strain this level was about 7% (Figure 4C). However, upon miltefosine addition, the wild-type and the Δ*rmiA::rmiA^+^* germlings were about 12% stained by PI (Figure 4C).,while more than 50% of the Δ*rmiA* germlings showed PI staining (Figure 4C). These results suggest that miltefosine induces both mitochondrial fragmentation and necrosis cell death in *A. fumigatus.*, which was accentuated in Δ*rmiA*, emphasizing the importance of RmiA for survival and viability of *A. fumigatus*.

**Figure 4.**
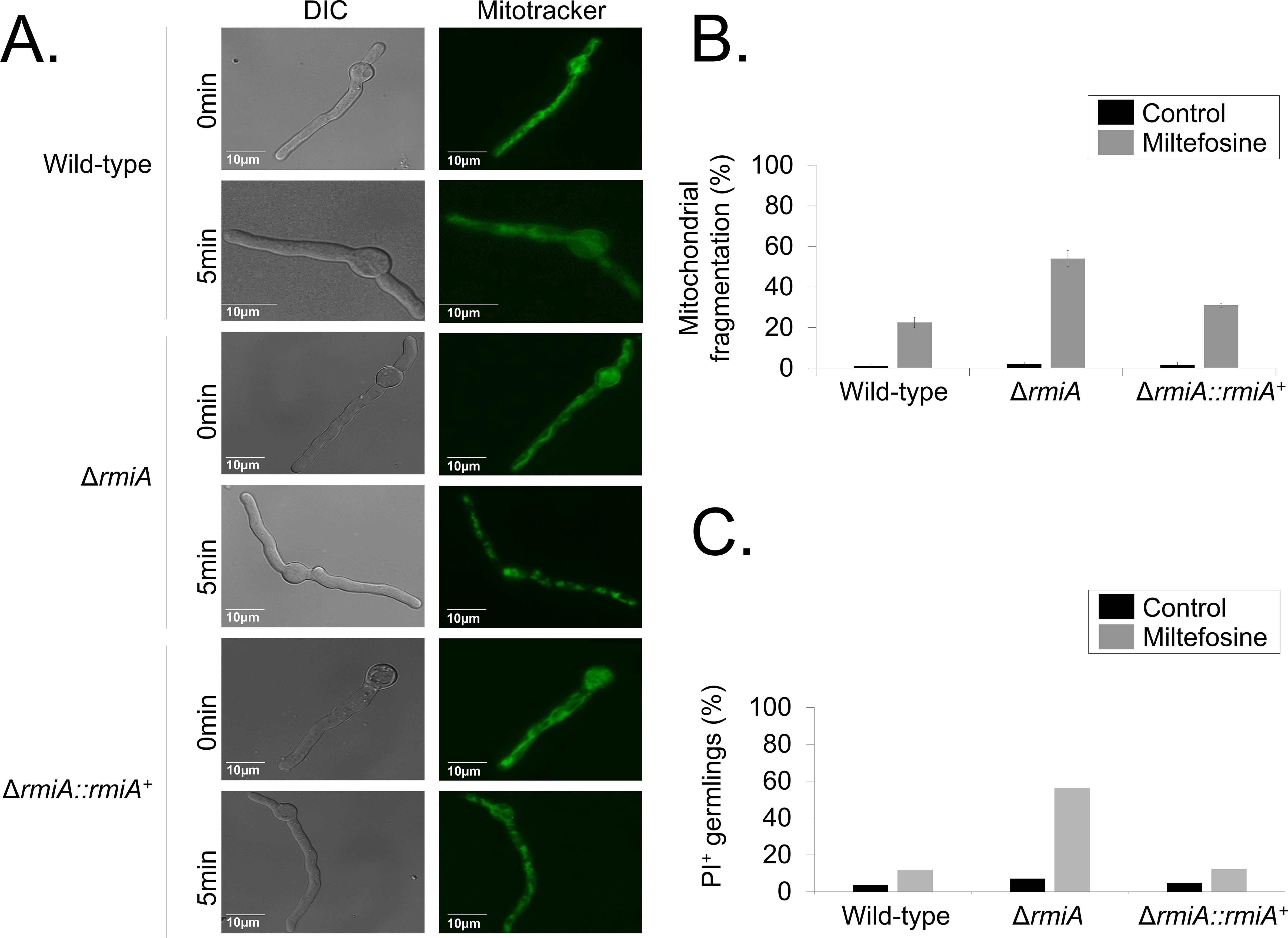
There is increased mitochondrial fragmentation and cell death when *A. fumigatus* Δ*rmiA* is exposed to miltefosine. (A) Mitochondrial morphology revealed by mitotracker in the wild-type, Δ*rmiA*, and Δ*rmiA::rmiA*^+^ strains. (B) Quantification of the mitochondrial fragmentation in the absence (control) and presence of miltefosine. (C) Quantification of PI^+^ (propidium iodide) germlings in the absence (control) and presence of miltefosine. The results are the average of three repetitions ± standard deviation of 30 germlings for each repetition.

*A. fumigatus* germlings were exposed to 4 µg/ml of a functional fluorescent analogue of miltefosine, MT-11C-BDP [11-(4,4-Difluoro-1,3,5,7-tetrametil-4-bora-3a,4a-diaza-s-indacen-2-il) n-undecilfosfatidilcolina] (86) for about 5 minutes (Figure 5). MT-11C-BDP localizes to tubular structures that resemble mitochondrial networks and that were also fragmented in a fraction of the germlings (Figures 5A and 5B). Colocalization with MitoTracker^TM^ Deep Red FM indicated that MT-11C-BDP analogue is mainly localized at the mitochondria.

**Figure 5.**
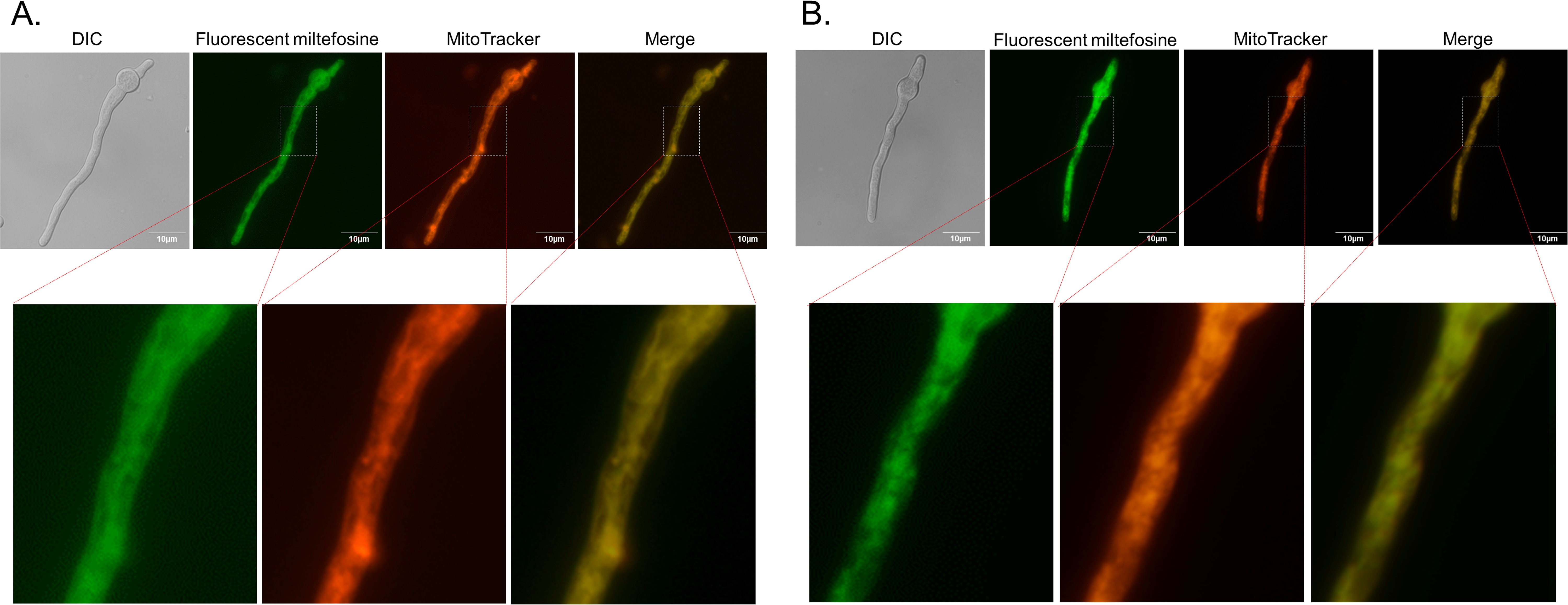
The fluorescent miltefosine analogue. MT-11C-BDP is localized in the mitochondria. (A) and (B) *A. fumigatus* germlings (16 h growth in MM) were exposed for 5 minutes to µg/mL of MT-11C-BDP. Germlings were stained with MitoTracker Deep Red^TM^ FM.

### Miltefosine induces the modulation of genes encoding proteins responsible for the metabolism of lipids, fatty acid and derivatives

Aiming to get insights about genes that are modulated under miltefosine exposure, we carried out a transcriptomic analysis (RNA-seq) comparing *A. fumigatus* wild-type strain exposed to miltefosine. In comparison to the wild-type grown in MM, when the cells were shifted to RPMI medium supplemented with 3 µg/ml of miltefosine for 30 minutes, a total of 1,248 genes were upregulated (log2FC >1.0; *P*<0.005) and 940 genes were downregulated (log2FC <-1.0; P<0.005). In both cases the false-discovery rate (FDR) was less than 0.05 (Supplementary Table S1).

The enrichment analysis using FunCat (https://elbe.hki-jena.de/fungifun/fungifun.php) showed a transcriptional upregulation of genes involved in vesicular and vacuolar transport, metabolism of glutamate, caspase activation, ABC transporters, osmosensing response, transport ATPases, stress response, proteasomal degradation, lipid transport and a high enrichment in lipid, fatty acid, and isoprenoid metabolism (Figure 6A). Genes involved in nuclear transport, RNA transport, Mitochondrial transport, TCA cycle, nucleotide binding, unfolded protein response, aminoacyl-tRNA-synthetases, amino acid metabolism, rRNA processing, ribosome biogenesis and translation were downregulated upon miltefosine exposure (Figure 6A). These results suggest that under miltefosine treatment, *A. fumigatus* increases the expression of genes involved in fatty acid metabolism and transport, stress responses and specific transporters while represses mitochondrial functions (e.g. TCA cycle and mitochondrial transport) and amino acid and protein biosynthesis.

**Figure 6.**
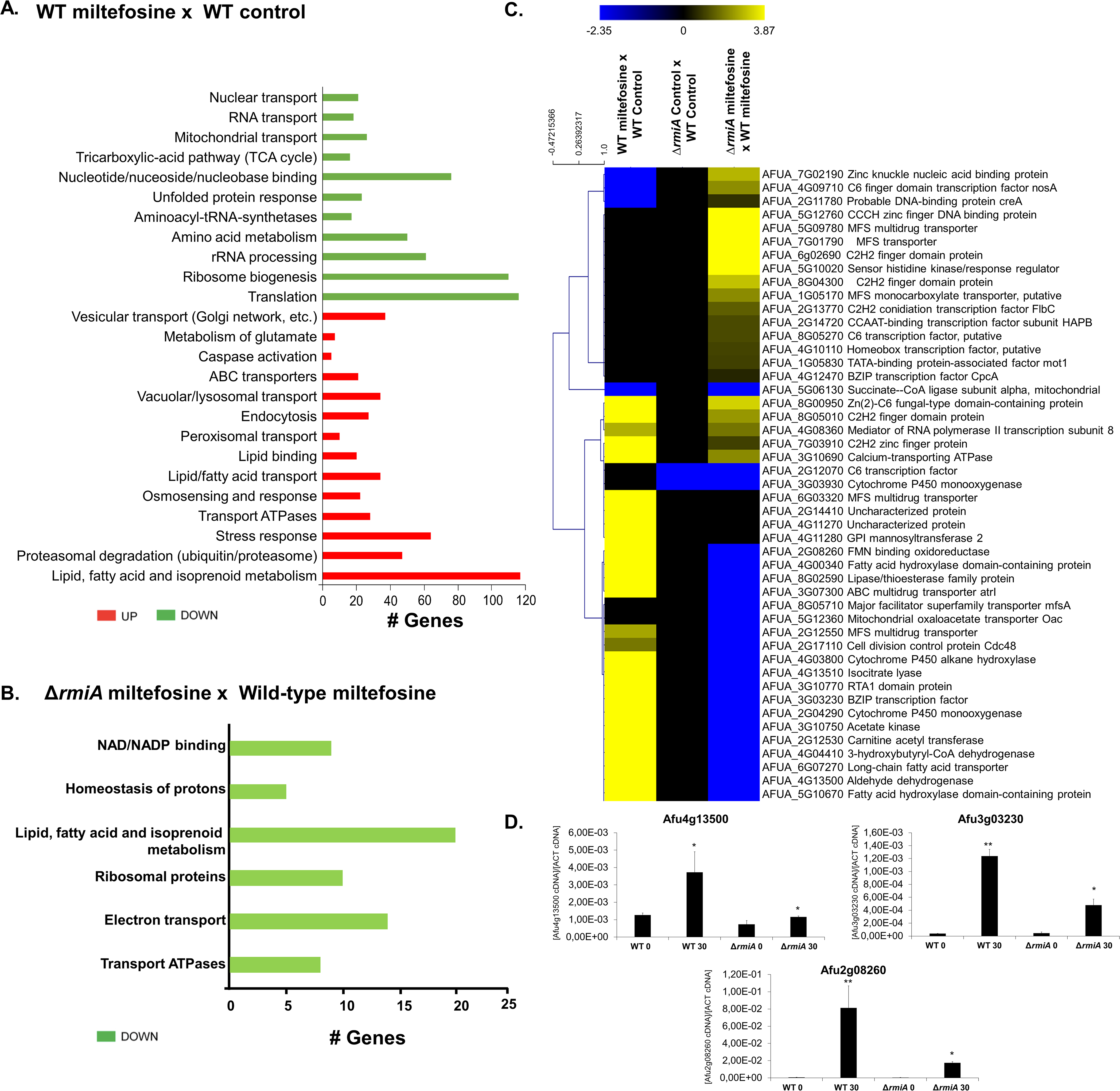
Transcriptional profiling of *A. fumigatus* wild-type and *ΔrmiA* exposed to miltefosine. (A) FunCat categorization of differently expressed genes (DEGs) up- and downregulated in the wild-type strain exposed to miltefosine in comparison to the wild-type strain grown in VMM (control). (B) FunCat analysis of DEGs downregulated in the Δ*rmiA* strain under miltefosine exposure in comparison to the Δ*rmiA* grown in VMM. (C) Heat map of log2 fold change (Log2FC) of DEGs as determined by RNAseq. Log_2_FC values are based on comparisons between (i) wild-type strain exposed to miltefosine *versus* wild-type strain in grown in MM (control); (ii) Δ*rmiA* grown in VMM (control) *versus* wild-type grown in VMM (control); (iii)) Δ*rmiA* strain exposed to miltefosine *versus* wild-type strain under miltefosine treatment. Hierarchical clustering was performed in Multiple Experiment Viewer (MeV) (http://mev.tm4.org/), using Pearson correlation with complete linkage clustering. Heat map scale and gene identities are shown. (D) Validation of RNA-seq data. Expression of three genes as determined by qRT-PCR after 0 and 30 minutes of exposition to 3µg/mL of miltefosine. Gene expression values were normalized by the expression of β-tubulin. Standard deviations are shown for biological triplicates.

### The *rmiA* is important for the induction of genes involved in lipid metabolism upon miltefosine exposure in *A. fumigatus*

To identify potential targets modulated by RmiA we performed the transcriptional profiling of the Δ*rmiA* null mutant under the same experimental design previously described for the wild-type strain. We identified 292 differentially expressed genes (DEGs), with 184 genes upregulated (log2FC >1.0; *P*<0.005) and 108 genes downregulated (log2FC <-1.0; P<0.005; FDR<0.05), upon miltefosine exposure (Supplementary Table S2). FunCat enrichment analysis has not shown categories for upregulated genes in the Δ*rmiA* mutant. However, FunCat for the downregulated genes in the Δ*rmiA* mutant exposed to miltefosine revealed enrichment for categories of genes encoding proteins involved in lipid, fatty acid and isoprenoid metabolism, NAD/NADP binding, homeostasis of protons, ribosomal proteins, electron transport and transport ATPases (Figure 6B).

A visual inspection of DEGs in both wild-type and Δ*rmiA* strains showed zinc finger proteins (AFUA_5G12760 and AFUA_6g02690), MFS transporters (AFUA_5G09780 and AFUA_7G01790) and a sensor histidine kinase regulator (AFUA_5G10020) with higher levels of expression in the Δ*rmiA* in comparison to the wild-type (Figure 6C). Genes involved in the metabolism of fatty acids (AFUA_4G00340, AFUA_2G12530, AFUA_6G07270, AFUA_5G10670, AFUA_8g02590), cytochrome P450 enzymes (AFUA_3g03930, AFUA_4G03800, AFUA_2G04290), cell division control protein (AFUA_2G17110), aldehyde dehydrogenase (AFUA_4G13500), isocitrate lyase (AFUA_4G13510) were downregulated in the Δ*rmiA* when compared to the wild-type strain (Figure 6C). Accordingly, the RNAseq data was validated by performing real-time PCR on 3 selected genes that showed a very similar expression pattern in comparison with data from the RNAseq (Figure 6D).

Taken together our data shows that in the wild-type strain, the lipid and fatty acid metabolism are upregulated upon miltefosine exposure suggesting their possible importance for survival in presence of this drug. On the other hand, the deletion of *rmiA* gene leads to a deficiency in the lipid and fatty acid metabolism, strongly suggesting that it could be linked to the higher sensitivity of this mutant to miltefosine.

### RmiA binds to a discrete number of gene promoter regions specifically in the presence of miltefosine

Considering that RmiA seems to be a TF involved in miltefosine resistance in *A. fumigatus*, we decided to identify potential direct targets under RmiA control of this protein using the Chromatin Immunoprecipitation coupled to DNA sequencing (ChIP-seq) approach. The RmiA:3xHA strain (Supplementary Figure S2 and Figure 7A) was grown in MM and further exposed to RPMI supplemented (or not) with miltefosine for 30 min. After immune-precipitation using anti-HA antibody, samples were sequenced using the Illumina HiSeq2500 platform, the reads were aligned to the A. *fumigatus* Af293 reference strain and the program MACS2 was used for peak calling. The peak intensity map showed that RmiA binding was enriched at the promoter region of 12 specific genes which are present in different chromosomes: (i) AFUA_2G08260 encoding a homologue of *S. cerevisiae* Oye2p, a NADPH oxidoreductase containing flavin mononucleotide (FMN) that may be involved in sterol metabolism, oxidative stress response, and programmed cell death (www.yeastgenome.org), (ii) AFUA_3G03230 encoding a BZIP transcription factor (www.fungidb.org), (iii) AFUA_3G07300 AtrA encoding an ABC multidrug transporter (www.fungidb.org), (iv) AFUA_3G10770 encoding a homologue of *S. cerevisiae* Rbs1p, a sphingoid long-chain base (LCB) efflux transporter; integral membrane transporter that localizes to the plasma membrane and may transport long chain bases (LCBs) from the cytoplasmic side toward the extracytoplasmic side of the membrane; role in glycerophospholipid translocation (www.yeastgenome.org), (v) AFUA_4G03800 encoding a cytochrome P450 alkane hydroxylase (www.fungidb.org), (vi) AFUA_4G13500 encoding a homologue of *S. cerevisiae* Hfd1p, a dehydrogenase involved in ubiquinone and sphingolipid metabolism, converting hexadecenal to hexadecenoic acid in sphingosine 1-phosphate catabolism, the human homologue of ALDH3A2, mutated in Sjogren-Larsson syndrome (www.yeasgenome.org and (87), (vii) AFUA_5G10670 encoding a protein that has domain(s) with predicted iron ion binding, oxidoreductase activity and role in fatty acid biosynthetic process, oxidation-reduction process (www.fungidb.org), (viii) AFUA_2G14410 encoding an orthologue that has a role in xanthophyll metabolic process (www.fungidb.org), (ix) AFUA_4G11270 encoding an unknown function hypothetical protein (www.fungidb.org), (x) AFUA_4G11280 encoding an orthologue that has dolichyl-phosphate-mannose-glycolipid alpha-mannosyltransferase activity and role in GPI anchor biosynthetic process (www.fungidb.org), (xi) AFUA_5G10660 encoding a pentatricopeptide repeat protein (www.fungidb.org), and (xii) AFUA_6G03320 encoding a MFS transporter (www.fungidb.org) (Figure 7A and Supplementary Table S3). The RmiA binding to these promoter regions happens specifically in presence of miltefosine suggesting that RmiA is important for the activity of those genes in the presence of this drug. Accordingly, the RNAseq data demonstrate that the expression levels of AFUA_2G08260, AFUA_3G03230, AFUA_3G07300, AFUA_3G10770, AFUA_4G03800, AFUA_4G13500, and AFUA_5G10670 are repressed in the Δ*rmiA* strain in comparison with the wild-type when both strains are exposed to miltefosine (Figure 7B and Supplementary Table S3).

**Figure 7.**
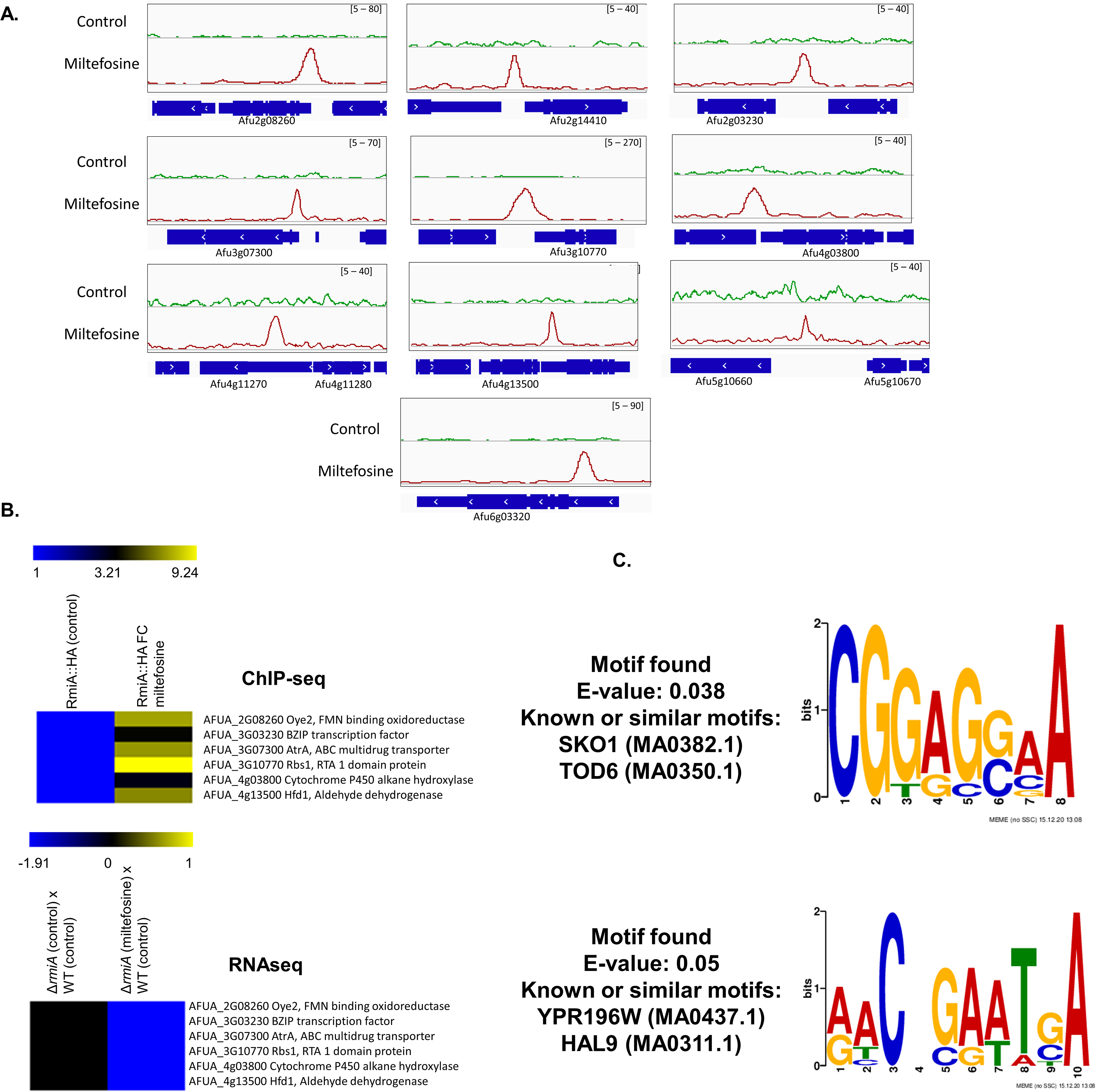
Chromatin Immunoprecipitation coupled to next generation sequencing (ChIP-Seq) of the wild-type and the Δ*rmiA* strains exposed to miltefosine. (A) ChIP-Seq Integrative Genomics Viewer (IGV; http://software.broadinstitute.org/software/igv/download) screenshot for promoter regions of genes that bound to RmiA:3xHA when grown for 24 h in VMM or after the addition of 12.5 µg/mL miltefosine for 30 min. (B) Heat map of the ChIP-seq results for 6 genes showing the fold enrichment of RmiA binding after 0 and 30 minutes of exposure to 12.5 µg/mL of miltefosine and the RNAseq results for the same 6 genes in the WT and Δ*rmiA* strains after exposure to 3µg/mL during 30 minutes. (C) MEME (Multiple EM for Motif Elicitation)-ChIP analysis of the 500 bp region surrounding the peaks identified in the ChIP-seq analysis.

To identify putative RmiA-binding motifs in *A. fumigatus,* we carried out Multiple Expectation maximum for Motif Elicitation (MEME) of the 500 nucleotides surrounding each peak sequence identified in the ChIP-seq. The results show the enrichment of two consensus DNA binding sequences for RmiA in the presence of miltefosine (Figure 7C). Two binding mofifs were predicted: 5’-CGGAG(G or C)AA-3’ (e-value of 5e-02) and 5’-AACNGAATGA-3’ (e-value of 3.8e-02).

Together, our data highlight the importance of RmiA for events involved with the miltefosine resistance process in *A. fumigatus* and suggest that genes potentially modulated by the RmiA binding have specific binding motifs for this protein. Several promoter regions of genes that are bound by RmiA encode proteins involved in lipid metabolism.

### RmiA is important for sphingolipids biosynthesis

Our previous results suggest that myriocin, a sphingolipid (SL) inhibitor, impairs the antifungal activity of miltefosine (Figure 1F). Considering that the metabolism of lipids seems to be involved with miltefosine resistance and the TF RmiA is linked to this process, we performed the SLs profiling of both wild-type and Δ*rmiA* strains exposed to miltefosine. Both strains were grown in VMM for 16h and shifted to RPMI media supplemented (or not) with 3 μg/ml of miltefosine for 4h. The main SL intermediates starting from the branching point of the pathway (DHS; dihydrosphingosine) were then measured through mass spectrometry analysis and the results were expressed in fold increase or decrease compared to the control not exposed to miltefosine (Figure 8A).

**Figure 8.**
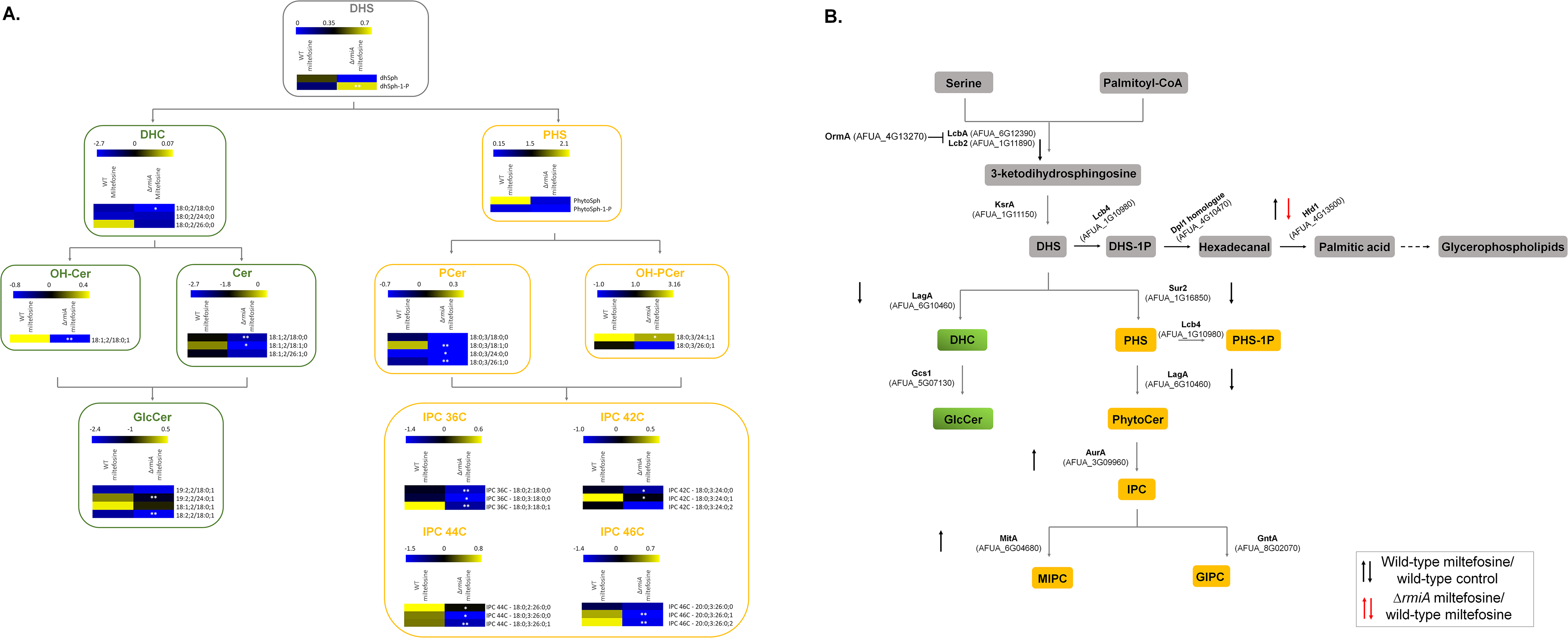
Deletion of *rmiA* leads to an overall reduction of sphingolipids biosynthesis in *A. fumigatus.* (A) The wild-type and Δ*rmiA* strains were grown in liquid VMM 16h and transferred to RPMI medium supplemented (or not) with 3 µg/mL miltefosine for additional 4h, the sphingolipids were extracted and measured by mass spectrometry. Heat maps labeled surrounded by boxes with gray borders represents the intermediates of SL biosynthetic pathwayheat maps surrounded by boxes with green borders represents the intermediates from the neutral branching while heat maps surrounded by boxes with yellow borders represents the intermediates from the acidic branch of SL biosynthetic pathway. Heat maps show the values obtained by the ratio VMM/RPMI. Experiments were performed by using three independent biological experiments and the results are the average of them. Statistical analysis was performed using Student’s t-test (p<0.05). (B) Diagram showing the different genes involved in the sphingolipids biosynthesis. Expression levels of genes encoding enzymes involved in the sphingolipids biosynthesis were selected from the RNAseq analysis. DHS, dihydrosphingosine; DHS-1P, dihydrosphingosine 1-phosphate; DHC, dihydroceramide; Cer, ceramide; GlcCer, glucosylceramide; OH-Cer, hydroxy-ceramide; PHS, phytosphingosine; PHS-1P, phytosphingosine 1-phosphate; PCer, phytoceramide; OH-PCer, hydroxy-phytoceramide; IPC, inositolphosphoryl ceramide.

The deletion of *rmiA* leads to an overall reduction of the analysed SLs when compared to the SL levels in the wild-type strain under the same conditions (Figure 8A). Interestingly, the reduction in SL levels occurs in both, acidic and neutral branches of the pathway, suggesting the deletion of *rmiA* affects the early steps of the SL biosynthetic process. The branching point of the SL pathway is DHS, the precursor of dihydroceramide (DHC; the first intermediary of the neutral branch) and the phytosphingosine (PHS; the first intermediary of the acidic branch). However, the DHS is also converted to dihydrosphingosine 1-phosphate (DHS-1P) starting the metabolic pathway where the DHS-1P is converted to glycerolipid through many enzymatic reactions (Figure 8A). Upon miltefosine exposure, there is an increase of DHS and a decrease in DHS-1P in the wild-type strain while the opposite is observed in the Δ*rmiA* mutant (Figure 8A). Phytosphingosine (PHS), ceramide (CER), hydroxyceramide (OH-CER), phosphoceramide (P-CER), hydroxyphosphoceramide (OH-PCER), glucoceramide (GLC-CER), and inositolphosphoryl-ceramide (IPC) are increased when the wild-type strain is exposed to miltefosine (Figure 8A). In contrast, all these sphingolipids were reduced in the Δ*rmiA* when exposed to miltefosine (Figure 8A). We investigated in our RNA seq dataset the expression levels of the genes that encode enzymes involved in the different steps of the SLs pathway (Figure 8B). We observed 5 genes (Lcb2, AFUA_1G11890; LagA, AFUA_6G10460; Sur2, AFUA_1G16850; and LagA, AFUA_6G10460) with reduced and 2 genes (AurA, AFUA_3G09960 and MitA, AFUA_6G04680) with increased expression when the wild-type was exposed to miltefosine (Figure 8B). Interestingly, only one gene (Hfd1, AFUA_4G13500) is differentially expressed with reduced expression when the Δ*rmiA* mutant is exposed to miltefosine (Figure 8B). Taken together, our results suggest that miltefosine antifungal activity against *A. fumigatus* interferes directly in the SL biosynthesis pathway.

### Azole-resistant clinical isolates of *A.* fumigatus are sensitive to miltefosine

to verify if miltefosine is a good candidate for therapy against azole resistant strains, we tested if miltefosine could inhibit *A. fumigatus* growth of 19 clinical isolates (in addition to CEA17 strain) with different levels of azole resistance by determining their MICs. We tested 9 azole-sensitive *A. fumigatus* strains (CEA17, CYP15-109, IF1S-F4, IFM59056, ISFT-021, IFM61407, MO68507, MO54056, and IFM59056) and also 10 azole-resistant isolates, with different resistance mechanisms, cultured from different sample sites from patients from Portugal, Japan, Belgium and Switzerland (88) (Table 2). The most common azole-resistance mechanisms include amino acid substitutions in the target Cyp51A protein and tandem repeat sequence insertions at the *cyp51A* promoter (54) The *cyp51A* gene is not mutated in the azole-resistant strains F16134, F14946, CYP15-117, CYP15-147, CYP15-75, CYP15-93, CYP15-106, and CYP15-115 (*cyp51A* was not sequenced in the CYP-15-91 strain), suggesting different mechanisms of azole resistance (88). In contrast, strains 1799392 and 20089320 have TR34 tandem repeats at the *cyp51A* promoter region and L98H amino acid replacement at the Cyp51A (89). All the azole-sensitive or –resistant clinical isolates have a MIC of 4 µg/ml of miltefosine (Table 2). These results strongly indicate that miltefosine can inhibit the growth of clinical isolates that have developed resistance to azoles through different mechanisms.

**Table 2.** Minimum inhibitory concentration (MIC, concentration in µg/µl) of *A. fumigatus* clinical isolates in the presence of different antifungals drugs

| Strains | Itraconazole | Posaconazole | Voriconazole | Amphotericin B | Miltefosine |
| --- | --- | --- | --- | --- | --- |
| CEA17 | 2 | 2 | 1 | 0.5 | 4 |
| F16134 | >8 | >8 | 4 | 0.25 | 4 |
| F14946 | >8 | 8 | 8 | 0.25 | 4 |
| CYP15-117 | >8 | 2 | 2 | 0.5 | 4 |
| CYP15-147 | >8 | 2 | 8 | 1 | 4 |
| 20089320 | >8 | 4 | 4 | 1 | 4 |
| CYP15-75 | 8 | 4 | 8 | 0.5 | 4 |
| CYP15-91 | 8 | 1 | 2 | 1 | 4 |
| CYP15-93 | 8 | 1 | 2 | 0.5 | 4 |
| CYP15-106 | 8 | 2 | 1 | 1 | 4 |
| CYP15-115 | 8 | 0.5 | 0.5 | 0.25 | 4 |
| 17993925 | 8 | 2 | 4 | 1 | 4 |
| CYP15-109 | 4 | 1 | >8 | 0.5 | 4 |
| IF1S-F4 | 4 | 1 | 1 | 0.5 | 4 |
| IFM59056 | 4 | 1 | 0.125 | 0.5 | 4 |
| ISFT-021 | 2 | 1 | 0.25 | 1 | 4 |
| IFM61407 | 2 | 2 | 0.125 | 0.5 | 4 |
| MO68507 | 1 | 1 | 0.25 | 0.5 | 4 |
| MO54056 | 1 | 1 | 0.125 | 0.5 | 4 |
| IFM59056 | 1 | 1 | 0.125 | 0.5 | 4 |

### Miltefosine increases the survival of *Galleria mellonella* larvae infected with *A. fumigatus*

On the basis of its essential role in sphingolipid biosynthesis, we asked whether RmiA is important for *A. fumigatus* virulence. *G. mellonella* larvae (n = 10 for each strain) were infected with the wild-type, *rmiA* deletion and complementation strains, and survival was assessed over a time period of 10 days (Figure 9A). The wild-type, Δ*rmiA*, and Δ*rmiA::rmiA^+^* strains caused 90 to 100 % mortality after 9 to 10 days postinfection (p.i.) (P < 0.001, Figure 9A). These results indicate that RmiA is not a key regulator of *A. fumigatus* pathogenesis in the *G. mellonella* model.

As a proof-of-principle of the *in vivo* antifungal activity of miltefosine, we tested its ability to control or reduce the *A. fumigatus* infection in *G. mellonella* larvae (Figures 9B and 9C). First, we tested three different concentrations of miltefosine (40, 60 and 80 mg/kg of the larva) aiming to verify the drug concentration that could cause minimal damage to the larvae. These miltefosine concentrations caused about 40 % mortality after 10 days (P < 0.001, Figure 9B). *A. fumigatus* infection of *G. mellonella* larvae combined with the different miltefosine concentrations resulted in about 50 % survival of the invertebrate host (P < 0.001, Figure 9C). These results indicate that miltefosine is able to control 50 % of the mortality caused by *A. fumigatus* infection in *G. mellonella*.

## Discussion

In the last years, the incidence of fungal infections has grown dramatically leading to an increasing number of deaths worldwide (1, 2). The mortality rate is linked to a set of conditions such as the host immune system integrity, availability of an effective antifungal drug, and the occurrence of clinical resistant isolates (6, 7, 11, 28, 90). The invasive pulmonary aspergillosis (IPA) is a disease caused by the opportunistic human pathogen *A. fumigatus* and displays high levels of morbidity and mortality mainly in immunocompromised patients (1, 24). Azoles are the main drug used to control IPA but the azole-resistant *A. fumigatus* isolates have increased significantly over the last decade (35, 40–45).

Given this scenario, there is an urgent need for new antifungal therapies applied to control IPA and other fungal diseases. The development of new antifungal drugs raises challenges such as the high costs and the time required for development and licensing of new compounds. To circumvent the slowness and cost of developing new drugs, the screening of chemical libraries and repurposing of drugs that are already commercialized for other purposes is a great opportunity to discover new antifungal compounds (62, 64, 67, 71, 91–93). Here we screened the growth of *A. fumigatus* in the presence of compounds present in two drug libraries and identified 10 compounds, among them five compounds already known as inhibitors of fungal growth including two azole-derivatives (econazole nitrate, and oxiconazole nitrate), fluvastatin, that inhibits ergosterol biosynthesis, and iodoquinol and miltefosine, drugs with an unknown mechanism of action. To our knowledge, the other five identified compounds (mesoridazine, cisapride, indinavir sulfate, enalaprilat, and vincristine sulfate) are novel as antifungal agents and have not been reported before. We investigated a possible mechanism of action for miltefosine, a chemical belonging to the alkylphosphocholine class. Miltefosine is mainly localized in the mitochondria and it has a MIC value of 4 µg/ml under *in vitro* conditions and we demonstrated that miltefosine is able to inhibit to the same extent *A. fumigatus* growth of several clinical isolates, including highly azole-resistant strains. Miltefosine was a drug initially used as an anti-neoplastic drug (94) and for treatment against trypanosomatids (95), and it is the first drug approved for oral treatment of leishmaniasis (77). However, the mechanism of action of miltefosine is not fully understood and not necessarily the same in different organisms, and the specific target of miltefosine has not been identified yet. Recent studies in trypanosomatids have suggested that miltefosine acts by: (i) altering the correct functionality of the sterol and sphingolipid metabolism (77, 78), (ii) inhibiting the phosphatidylcholine synthesis (96), and the membrane remodeling due to the phospholipase action, thus contributing to changing membrane physical properties (97), (iii) inhibiting cytochrome *c* oxidase (98), (iv) activating the plasma membrane Ca^+2^ channel opened by the sphingolipid sphingosine and (v) destabilizing the intracellular Ca^+2^ homeostasis (78). On the other hand, the resistance phenotype to miltefosine in trypanosomatids has been linked to genes belonging to lipid metabolism (99).

Concerning to its antifungal behavior, miltefosine has demonstrated to be effective against different fungal species (100–111), however, its mode of action remains to be clarified. Recent studies have suggested that miltefosine triggers its antifungal effects by destabilizing cell membranes and inducing apoptosis (102, 105, 110, 112, 113). Accordingly, Spadari and colleagues (2018) demonstrated that for *Cryptococcus spp.* miltefosine affects the plasma membrane permeability due its interaction with ergosterol and/or phospholipids increasing the production of ROS and DNA fragmentation which culminates with the fungal death by apoptosis (105). In addition, in *C. krusei,* the mode of action of miltefosine is also supposed to be related to the binding of the drug to ergosterol in the cell membrane leading to the cell apoptosis (110).

We observed that at the MIC concentration, miltefosine displayed a fungicidal effect against *A. fumigatus,* corroborating with previous results presented for several fungal species such as *Cryptococcus* spp, *Candida* spp, and molds (101–106, 108–110, 114). Our studies showed that miltefosine could decrease 50 % *A. fumigatus* mortality in *G. mellonella* larvae. We have then decided to check if miltefosine could present any interaction with other antifungal drugs. Azoles such as posaconazole and voriconazole that act by inhibiting the ergosterol biosynthesis (115), amphothericin B that sequesters ergosterol from the cell membrane (116) and caspofungin that targets the glucan synthase Fks1 and inhibits the synthesis of β-(1, 3) glucan (117) were included in our analysis and none of them showed interaction with miltefosine. Our results corroborated what was previously observed in *Aspergillus spp* where the interaction between these compounds with miltefosine was indifferent for 32 from 33 isolates (103). In contrast to our results, miltefosine has been reported to have synergy with posaconazole against *Fusarium oxysporum* and the mucormycetes (118). In addition, a recent study with *C. auris* demonstrated that for 25% of the isolates assessed there was a synergic activity between miltefosine and amphothericin B, with FICI=0.5 (110).

Sphingolipids (SL) are complex lipids composed of octadecacarbon alkaline blocks, synthesized from non-sphingolipid precursors, and representing one of the most abundant lipids in the eukaryotic cell membranes (119, 120). In fungi, SLs are involved in central cellular functions such as growth, pathogenesis, cell death, and signal transduction (121–123). The SL biosynthesis starts in the endoplasmic reticulum where the non-lipidic precursors serine and palmitoyl coenzyme A are condensed by the serine palmitoyltransferase enzyme (SPT) into 3-keto dihydrosphingosine. The SPT is specifically targeted by myriocin, a sphingolipid inhibitor (79). The interaction assay between miltefosine and myriocin showed that at high concentrations of both compounds, the FICI value was greater than 4.0 characterizing an antagonistic effect between these drugs (76). This implicates that sphingolipid metabolism may be important to the antifungal effect of miltefosine, corroborating with previous results obtained for other fungal species and trypanosomatids (77, 78, 105, 110).

We were able to identify a completely novel *A. fumigatus* transcription factor RmiA, linked to miltefosine resistance in this pathogen. This information came from a large-sale phenotypic screening of a collection of TF deletion mutants in presence of miltefosine. Although the deletion of six TFs somehow moderately impacted the growth of the mutant in presence of miltefosine, the Δ*rmiA* is the most sensitive mutant. RmiA is a novel and uncharacterized TF that codifies a putative Zn(II)Cys6 binuclear domain that translocates to the nucleus in presence of miltefosine and seems to be a key TF in the miltefosine response in *A. fumigatus*. Miltefosine at MIC completely abolished the Δ*rmiA* mutant growth and no additional phenotypes were observed under other stress conditions such as growth in presence of sub-inhibitory concentrations of posaconazole, voriconazole, caspofungin, NaCl, Calcofluor white, sorbitol, and CaCl_2._ The Identification of the *rmiA* gene as a putative major TF involved in *A. fumigatus* response to miltefosine provided us with an opportunity to enquiry the molecular mechanisms that are regulated by this gene.

The transcriptional profiling through RNAseq assay with the wild-type strain in presence or absence of miltefosine indicated increased upregulation of genes involved in lipids/fatty acid transport and metabolism. In contrast, the RNAseq of the Δ*rmiA* mutant exposed to miltefosine shows exactly an opposite behavior. The lipid and fatty acid metabolism were the main category of down-regulated genes which strongly suggests that this TF participates directly or indirectly in the induction of genes involved in lipid metabolism, including genes involved in the biosynthesis of sphingolipids. The identification of RmiA represents the first genetic element described and characterized which plays a direct role in miltefosine response in fungi.

Our work provides opportunities for understanding the mechanism of action of miltefosine throughout the characterization of the genes that are differentially expressed in the Δ*rmiA* mutant. Further work will focus on the molecular characterization of these differentially expressed genes.

## Materials and Methods

### Media, strains and phenotypic characterization

The *Aspergillus spp.* used in this work are listed in the Table S1. Azole-resistant *Aspergillus fumigatus* strains were kindly provided by Katrien Lagrou and were isolated from different sources in Belgium, Switzerland, Portugal, EUA. All *Aspergillus* strains were grown in either solid minimal medium [MM: 1 % glucose (w/v), 50 mL of a 20× salt solution, trace elements, 2% agar (w/v) pH6.5] or solid complete medium [YAG: 2% glucose (w/v), 0.5% yeast extract(w/v), trace elements, 2% agar (w/v)] at 37°C. The composition of the trace elements and nitrate salts is described at Käfer, 1977. For RNAseq, ChIP-seq and lipidomics, conidia were germinated in RPMI-1640 media and transferred to liquid Vogel’s Minimal Medium (VMM). For phenotypic characterization, plates containing solid MM were centrally inoculated with 10^5^ spores of each strain in presence or absence of variable concentration of miltefosine [0-8µg/mL]. After 120h of incubation at 37°C, the radial growth was measured. All plates were grown in triplicate, and averages ± standard deviations (SD) of the data are plotted. All strains used in this work are listed in the Supplementary Table S4.

### Library drug screenings

Two different drug libraries were screened for antifungal activity against *A. fumigatus* CEA17 strain: The Pathogen Box (https://www.mmv.org/mmv-open/pathogen-box) and the National Institutes of Health (NIH) clinical collection (NCC) (https://pubchem.ncbi.nlm.nih.gov/source/NIH%20Clinical%20Collection). The Pathogen Box (https://www.mmv.org/) is a collection of 400 diverse, drug-like molecules with already described activity against different pathogens responsible for important neglected diseases such as malaria, tuberculosis, toxoplasmosis, and others. The NCC library is composed by a small molecule repository of 727 compounds which are part of screening library for the NIH Roadmap Molecular Libraries Screening Centers Network (MLSCN) corresponding to a collection of chemically diverse compounds that have been in phase I-III clinical trials (64).

For the primary screening, the drugs were diluted from 0.78-25µM in 200µL of MOPS [3-(N-morpholino) propanesulfonic acid] buffered RPMI 1640 (Life Technologies), pH 7, in 96-well plates. In each well a total of 1×10^4^ conidia of *A. fumigatus* wild-type strain was inoculated. Plates were incubated for 48h at 37°C without shaking. Wells containing only medium and DMSO were used as controls. Fungal growth inhibition was determined visually as a no-growth endpoint and those compounds were selected for further studies. All experiments were done in triplicate.

Fungicidal or fungistatic activity of the selected compounds was also assessed. Briefly, a total of 1×10^4^ conidia of *A. fumigatus* wild-type strain was inoculated in 96-wells plates, each well containing 200µL of MOPS buffered RMPI 1640 medium plus the lowest concentration of each compound that promoted the fungal growth inhibition in the primary screening. Plates were incubated 48h at 37°C without shaking. Following, 100 was plated in solid complete medium and incubated at 37°C for another 36h. Wells containing only medium and DMSO were used as controls. The number of viable colonies was determined by colony-forming unity (CFU) compared to the negative control (no drug), which had 100% survival. Results are expressed as means standard deviations (SD) of three independent experiments.

### Minimal inhibitory concentration (MIC)

The miltefosine drug used for minimal inhibitory concentration (MIC) assays was purchased from Sigma-Aldrich and solubilized in ethanol. The MIC was determined based on the M38-A2 protocol of the Clinical and Laboratory Standards Institute (125).

Briefly the assay was performed in 96-wells plates containing 200µL of MOPS buffered RPMI 1640 medium pH 7.0, supplemented with miltefosine [0-8µg/mL] and 1×10^4^ conidia of *A. fumigatus,* per well. Plates were incubated at 37°C without shaking for 48h. Wells containing only medium and ethanol were used as control. The MIC value was defined as the lowest concentration of miltefosine that visually inhibited 100% of fungal growth. All experiments were done in triplicate.

### Assays for checking antifungal activity of drug combinations

We checked the interaction of miltefosine with several drugs including antifungals and lipid inhibitor using a checkerboard microdilution method. The drugs concentrations ranged from 0.001 to 8.0 µg/mL for miltefosine, 0.03 - 2.0 µg/mL for posaconazole, 0.0007 - 0.5 µg/mL for voriconazole, 4.0 - 256.0 µg/mL for caspofungin and 0.06 - 4.0 µg/mL for amphotericin B and 2.0 – 128 µg/mL for myriocin. The plates were incubated at 37°C during 48h. The MIC endpoint was 100% of growth inhibition. The interaction was quantitatively evaluated by determining the fractional inhibitory concentration index (FICI): FICI = [MIC miltefosine in combination/MIC miltefosine] + [MIC clinical drug in combination/MIC clinical drug]. The FICI was calculated for all possible combinations of different concentrations (126). Also, interaction curves were constructed. The interaction between these drugs was classified as synergic if FICI ≤0.5, indifferent if 0.5 <FICI≤4.0, and antagonic for FICI >4.0 (76).

### Construction of *A. fumigatus* mutants

To generate the RmiA:3xHA mutant, a 2.9Kb fragment encompassing the rmiA ORF and the 5’UTR region, along with the 1kb 3’UTR DNA sequence, were PCR amplified from CEA17 genomic DNA (gDNA) with primer pairs P1/P2 and P4/P5 respectively. The 0.8kb 3xHA-trpC fragment was amplified from the pOB430 plasmid with primers P10/P11, the prtA gene was amplified from the plasmid pPTRI with primers P8/P9.

The RmiA:GFP strain was constructed by the amplification of a 2.9Kb fragment encompassing the rmiA ORF and the 5’UTR region, along with the 1kb 3’UTR DNA sequence, were PCR amplified from CEA17 gDNA with primer pairs P1/P3 and P4/P5 respectively. The GFP-trpC fragment was amplified from the pOB435 plasmid with primers P10/P11, the prtA gene was amplified from the plasmid pPTRI with primers P8/P9.

The Δ*rmiA* strain was complemented generating the Δ*rmiA::rmiA^+^* lineage. Specifically, the fragment containing the 5’UTR plus the *rmiA* gene, along with the 1kb 3’UTR DNA sequence, were PCR amplified from CEA17 gDNA with primer pairs P1/P7 and P4/P5 respectively. In addition, these fragments were fused to the *ptrA* gene, which was previously PCR amplified from plasmid pPRTI (primers P8/P9).

All DNA cassettes (rmiA^+^::prtA, rmiA::GFP::prtA, and rmiA::3xHA::prtA) were constructed by *in vivo* homologous recombination by using *S. cerevisiae* (127). Briefly, the set of fragments of each of the constructions along with the plasmid pRS426 digested with BamHI/EcoRI were transformed into the *S*. *cerevisiae* SC9721 strain. Whole cassettes rmiA^+^::prtA, rmiA::GFP::prtA and rmiA::3xHA::prtA were transformed into the *ΔrmiA* strain, candidates were selected by resistance to pyrithiamine and further verified via western blot, reversal of miltefosine sensitivity phenotype and/or protein functionality.

Primers used in this work are listed in the Supplementary Table S5. Additionally, the mutant strains constructed in the current work were performed into the background of the Δ*rmiA* strain. Positive candidates were selected in the presence of pyrithiamine, purified through three rounds of growth on plates, submitted to gDNA extraction, and confirmed by PCR.

### Protein extraction and immunoblot analysis

A total of 1 x 10^6^ conidia/mL of each strain was inoculated in 50 mL of Vogel’s medium and grown at 37 °C for 16h under agitation. Mycelia were then washed with RPMI 1640 media and incubated in RPMI 1640 containing 12.5 μg/mL of miltefosine at 37°C for 0, 4 and 8h at 37°C with shaking. For protein extraction, mycelia were ground into liquid nitrogen and resuspended in 0.5mL of lysis buffer (10% (v/v) glycerol, 50 mM Tris-HCl pH 7.5, 1% (v/v) Triton X-100, 150 mM NaCl, 0.1% (w/v) SDS, 5 mM EDTA, 50 mM sodium fluoride, 5 mM sodium pyrophosphate, 50 mM - glycerophosphate, 5 mM sodium orthovanadate, 1 mM phenylmethylsulfonyl fluoride (PMSF), and 1x complete mini protease inhibitor (Roche Applied Science). Extracts were centrifuged at 16,000 g for 20 min at 4°C. The supernatants were collected, and the protein concentrations were determined using the Bradford assay (Bio-Rad). Then, 30µg of total protein extract from each sample were resolved in 10% (w/v) SDS-PAGE and transferred to a nitrocellulose membrane for a Western blot assay. Monoclonal anti-HA antibody (Sigma-aldrich) was used to confirm the RmiA::3xHA expression. In addition, anti-α-actin antibody was used to normalize the protein loading. The primary antibodies were detected using a horseradish peroxidase (HRP)-conjugated secondary antibody raised in mouse (Sigma-Aldrich). Chemiluminescent detection was achieved using an ECL Prime Western blotting detection kit (GE Healthcare). To detect these signals on blotted membranes, the ECL Prime Western blotting detection system (GE Healthcare, Little Chalfont, UK) and LAS1000 (Fujifilm, Tokyo, Japan) were used.

### Real-time PCR analysis

Total cellular RNA was extracted using TRIzol reagent (Invitrogen, Life Technologies, Camarillo, CA, USA). Further, RNA was submitted to DNA digestion with RQ1 RNase-free DNase (Promega, Fitchburg, WI, USA) according to the manufacturer’s instructions. The cDNA synthesis was performed by the ImProm-II™ Reverse Transcription System (Promega) and oligo(dT). The real-time OCR was performed using the ABI 7500 Fast Real-Time PCR System (Applied Biosystems, Foster City, CA, USA) and the SYBR Green PCR Master Mix kit (Applied Biosystems), according to manufacturer’s instructions. Analysis were carried-out using three independent biological replicates. The mRNA quantity relative fold change data was calculated using standard curves (128) and normalized by the expression levels of the housekeeping β-tubulin gene. Primer sequences used in this study are listed in Supplementary Table S5.

### RNA purification and preparation for RNA-Seq

A total of 10^6^ spores/ml of *A. fumigatus* WT and Δ*rmiA* were inoculated in 50 ml of Vogel’s medium and grown at 37 °C for 16h under agitation. The suspensions were centrifuged and washed with PBS. Mycelia was suspended in VMM media containing glucose supplemented with 3 μg/mL of miltefosine, or in the absence of any drug, and incubated at 37 °C for additional 30 min. Total RNA was extracted by Trizol method. Subsequently, 10 µg of total RNAs were subjected to RNA purification using DNase I (New England Biolabs Inc.) and the quality was checked on 2% agarose gel and verified using the Agilent Bioanalyser 2100 (Agilent technologies). RNAs selected for further analysis had a minimum RNA Integrity Number (RIN) value of 8.0. One µg of purified RNAs were used for library preparation using Illumina NEBNext® Ultra^TM^ Directional RNA Library Prep Kit according to manufacturer’s protocol and sequenced using the Illumina HiSeq2500 platform at the Genomics and Single Cells Analysis Core facility at the University of Macau. The expression levels were calculated in RPKM and for the differential expression analysis a log2FoldChange of -1≤Log2FC≥1 was applied to capture minimum 2 times perturbation on the expression levels, with the P<0.005 and false-discovery rate (FDR) lower than 0.05.

### Chromatin preparation

A similar experimental design used for the RNA seq was used for the ChIP seq experiments. After growth, the cultures were added with 1% formaldehyde for 20 minutes with gentle shaking at room temperature, then a final concentration of 0.5M glycine was added to further incubation for 10 minutes. The mycelia were collected by filtering and washed with cold water. The crosslinked mycelia were frozen in liquid nitrogen and frozen dried for 2 hours before lysis. The cell lysis was processed by 6 times beating for 3 minutes with ∼100 μl volume of silica beads using Bullet Blender (Next Advance) with 3 minutes of cooling in between each cycle. Chromatins were extracted as described (129) and sonicated using the Qsonica Q800R at 100% amplitude with 10 seconds ON and 15 seconds OFF cycles for a total sonication time of 30 minutes. Chromatin concentration and size (100-500bp) were checked on 2% agarose gel, and the prepared chromatins were stored at −80 ℃ until use.

### Chromatin Immuno-precipitation and sequencing library preparation

Immuno-precipitation was carried out using anti-HA antibody as described previously (130). Immuno-precipitated materials were purified using QIAGEN PCR clean-up kit, and multiplexed sequencing libraries were prepared as described (130) using NEBNext® Ultra™ II DNA Library Prep Kit for Illumina according to manufacturer’s protocol. Libraries were checked and quantified using DNA High Sensitivity Bioanalyser assay, mixed in equal molar ratio and sequenced using the Illumina HiSeq2500 platform at the Genomics and Single Cells Analysis Core facility at the University of Macau.

### Data mapping and bioinformatics analysis

Raw sequencing reads of ChIPseq experiments were quality-checked using FastQC (http://www.bioinformatics.babraham.ac.uk/projects/fastqc/) and aligned to Af293 reference genome (genome version s03-m05-r06) using Bowtie2 (version: 2.2.9) (131). For peaks calling, MACS2 was applied. In order to determine the presence of conserved RmiA DNA binding motifs we carried out a MEME (Multiple EM for Motif Elicitation)-ChIP analysis to search the 500 bp region surrounding the peaks identified in our ChIP-seq data (http://meme-suite.org).

### Lipid analysis

A total of 10^6^ spores/ml of *A. fumigatus* WT and ΔAfub_027770 were inoculated in 50 ml of Vogel’s medium and grown at 37 °C for 16h under agitation. The suspensions were centrifuged and washed with PBS. Mycelia was suspended in VMM media containing glucose supplemented with 3 μg/mL of miltefosine, or in the absence of any drug, and incubated at 37 °C for additional 4h. Prior to cell lysis, C17-sphingolipids were added to the samples (132, 133). Mandala extraction was carried out as described previously (134), with a few modifications. To facilitate the disruption of mycelia, the samples were vortexed and sonicated for 2 min in the presence of 0.2 g of glass beads. Then, the samples were submitted to Bligh and Dyer Extraction (135). A quarter of each sample obtained from the Bligh and Dyer Extraction was reserved for inorganic phosphate (Pi) determination, so the relative sphingolipid signal was normalized by the Pi abundance. The organic phase was transferred to a new tube and submitted to alkaline hydrolysis of phospholipids (136). Finally, the organic phase was dried and used for mass spectrometry analysis (133).

### Statistical analysis

Grouped column plots with standard deviation error bars were used for representations of data. For comparisons with data from wild-type or control conditions, we performed one-tailed, paired *t* tests or one-way analysis of variance (ANOVA). All statistical analysis and graphics building were performed by using GraphPad Prism 5.00 (GraphPad Software).

### Fluorescence microscopy

A total of 10^5^ spores of each strain was inoculated on coverslips in 4 ml of MM medium for 16 h at 30°C. After, coverslips with adherent germlings were left untreated or miltefosine for different periods of time, as indicated. Staining procedures included: (i) 5 min of incubation in a solution with Propidium Iodide (PI [0.05mg/ml], Sigma Aldrich); (ii) 5min of incubation in a solution with MitoTracker Deep Red^TM^ FM dye ([250nM], Invitrogen); (iii) 10min of incubation in a solution containing Hoechst 33342 dye (20µg/mL, Molecular Probes, Eugene, OR, USA). Further, the coverslips were rinsed with phosphate-buffered saline (PBS; 140 mM NaCl, 2 mM KCl, 10 mM NaHPO4, 1.8 mM KH_2_PO_4_, pH 7.4). Slides were visualized on the Observer Z1 fluorescence microscope using a 100x objective oil immersion lens. Differential interference contrast images (DIC) and fluorescent images were captured with AxioCam camera (Carl Zeiss) and processed using AxioVision software (version 4.8). In each experiment, at least 50 gemlings were counted. For GFP the wavelength excitation was 450 to 490 nm and emission wavelength of 500 to 550 nm. For MitoTracker Deep Red^TM^ FM the wavelength abs/em was about 644/665 nm. For Hoechst [4,6-diamidino-2-phenylindole] staining, the excitation wavelength was 365 nm and emission wavelength 420-470 nm. For PI the wavelength excitation was 572/25 nm and emission wavelength was 629/62 nm.

### Virulence analysis in *Galleria mellonella* model

The *Galleria mellonella* larvae were obtained by breeding adult larvae (137) weighting 275-330 mg. The larvae were kept in starvation in Petri dishes at 37° in the dark for 24 hours prior to infection. The larvae used for the experiment were in the sixth stage of development. For infection, fresh spores from each strain (mutants and wild-type) were used. The spores of each strain were counted using a hemocytometer. The stock concentration of spore suspensions used for infection was 2×10^8^ conidia/ml, and from this stock 5ul were used for larval infection (1×10^6^ conidia/larva). The control group was composed of larvae inoculated with 5 μl of PBS to observe any possible death caused by physical trauma. The inoculum was performed using a Hamilton syringe (7000.5KH) and the conidia were inoculated into the lower left proleg of the larvae. After 30 minutes of the larvae being infected, treatment with miltefosine (M5571 Sigma-Aldrich) was carried out. The drug was rehydrated in distilled water as recommended by the manufacturer (102). The concentrations used for the treatments were 40, 60 and 80 mg/kg of the larvae and each larva was weighted individually, and the volume was adjusted to the pre-established concentrations. As a control for the treatments, we made three groups of larvae in which concentrations of 40, 60 and 80mg / kg were injected. The treatments were also injected into the lower right proleg of the larvae. After infection, the larvae were kept at 37 ° in Petri dishes in the dark and scored daily. Larvae were considered dead due to lack of movement in response to touch. The viability of the inoculum administered was determined by plating a serial dilution of the conidia in YAG medium. The statistical significance of the comparative survival values was calculated using the log rank analysis of Mantel-Cox and Gehan-Brestow-Wilcoxon by using the statistical analysis package Prism (138).

### Data availability statement

The datasets generated for this study are available on request to the corresponding author.

## Supplemental material

**Supplementary Figure S1.**
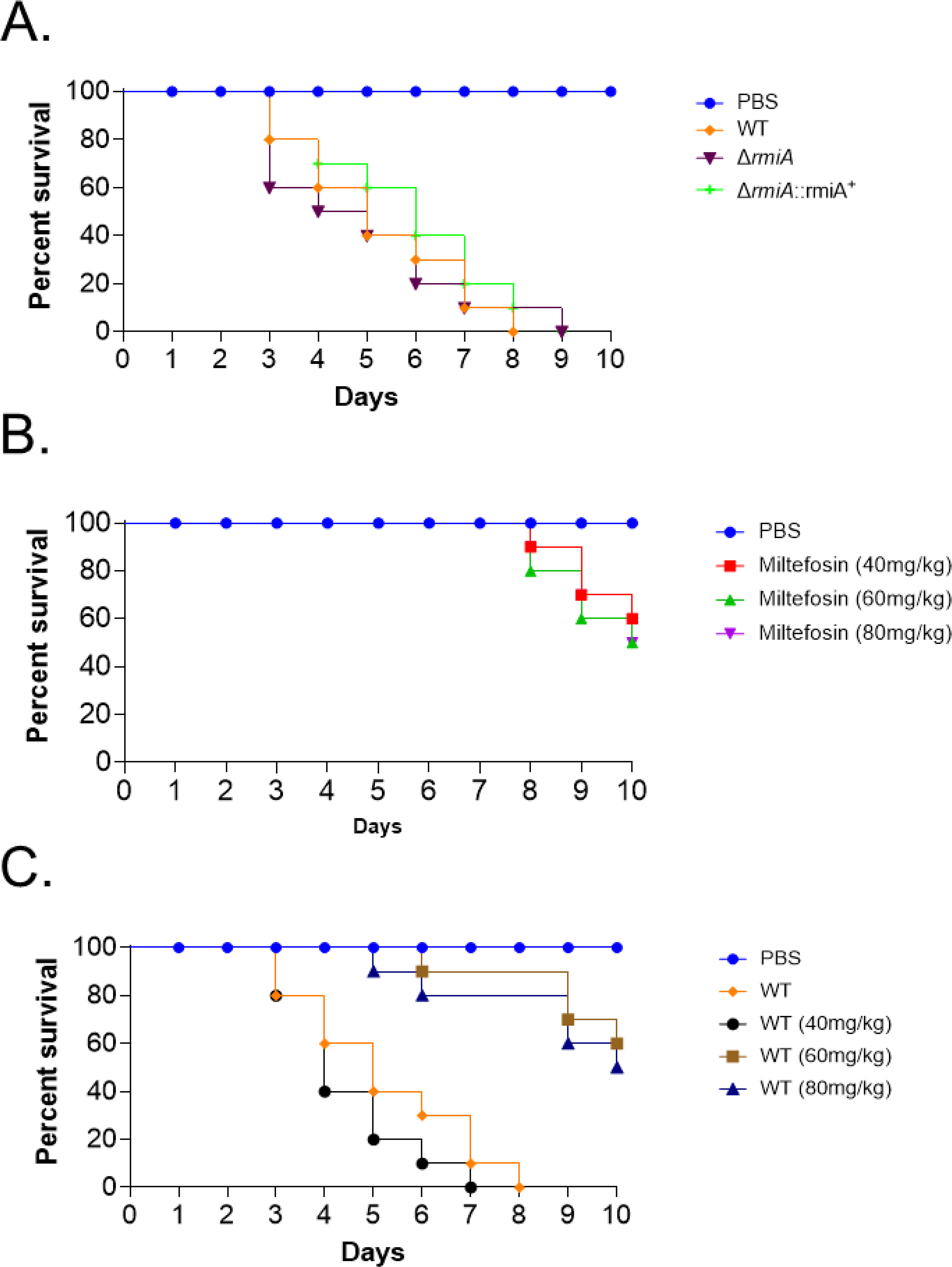
The deletion of *rmiA* causes no growth impact in the presence of different stress conditions. Strains were grown from 10^5^ spores for 5 days at 37°C on minimal media (MM) supplemented or not with (A) CaCl2, (B) calcofluor white (CFW), (C) sorbitol, (D) NaCl, (E) menadione and (F) variable temperature. The results are expressed as the average of radial diameter of the treatment divided by the radial diameter of the control of three independent experiments ± SD. Standard deviations present the average of three independent biological repetitions.

**Supplementary Figure S2.** Construction of *rmiA::3xHA* and *rmiA::GFP* strains. Both strains were constructed in the Δ*rmiA* background and confirmed by PCR sing *rmiA pRS426 3R* (P5) and *rmiA 1500UP ext F* (P6) primers. The *rmiA::3xHA* (A) and *rmiA::GFP* (B) are functional, do not present growth defects and restored the miltefosine sensitivity of the Δ*rmiA* null mutant.

**Supplementary Table S1.** Genes differentiallly expressed in the wild-type exposed to miltefosine.

**Supplementary Table S2.** Genes differentiallly expressed in the Δ*rmiA* mutant compared to the wild-type exposed to miltefosine.

**Supplementary Table S3.** Genes identified in the RmiA ChIP-seq.

**Supplementary Table S4.** Strains used in this work.

**Supplementary Table S5.** List of primers used in this work

